# SMG1:SMG8:SMG9-complex integrity maintains robustness of nonsense-mediated mRNA decay

**DOI:** 10.1101/2024.04.15.589496

**Authors:** Sabrina Kueckelmann, Sophie Theunissen, Jan-Wilm Lackmann, Marek Franitza, Kerstin Becker, Volker Boehm, Niels H. Gehring

## Abstract

Nonsense-mediated mRNA decay (NMD) is a translation-dependent mRNA turnover pathway, which degrades transcripts containing premature termination codons. SMG1-mediated phosphorylation of the key NMD factor UPF1 is essential for NMD initiation and regulated by SMG9 and the C-terminus of SMG8. However, their specific roles in NMD regulation within intact cells remain partially understood. Here, we deleted the C-terminus of endogenous SMG8 in human cultured cells, which resulted in unchanged NMD activity. Cell lines lacking SMG8 and SMG9 showed slight NMD inhibition and unchanged UPF1 phosphorylation levels, but were sensitized to treatment with a SMG1 inhibitor (SMG1i). Transcriptome-wide analysis revealed the upregulation of NMD-annotated transcripts, which corresponded to synergistic effects of SMG1i concentration and SMG8 and SMG9 knock-out conditions. Moreover, the UPF1 interactome showed enrichment of various NMD factors in SMG8 or SMG9 knock-out cells and following SMG1i treatment, suggesting an accumulation of stalled NMD complexes at various stages of the NMD process. Together, our work uncovers important roles of SMG8 and SMG9 in maintaining NMD robustness in human cells.

## Introduction

Alongside other co-translational quality control mechanisms, nonsense-mediated mRNA decay (NMD) detects and eliminates defective or undesired mRNAs ^1–3^. The primary role of NMD is to identify mRNAs carrying premature translation termination codons (PTCs) caused by mutations, alternative splicing, and other means ^4^. The activity of NMD prevents the production of C-terminally truncated and possibly toxic proteins ^5,6^. Beyond its role in quality control, NMD also plays a role in regulating gene expression, thereby directly or indirectly affecting approximately 20-40% of genes ^7^. In many cases, NMD is activated by the exon junction complex (EJC). This multi-protein, RNA-binding complex is deposited 20-24 nucleotides (nts) upstream of exon-exon junctions by the spliceosome and remains bound to the mRNA during its export to the cytoplasm ^8^. Translation termination at a PTC differs from that at a normal termination codon due to the presence of a downstream EJC, given that the PTC is positioned at least 50-55 nts upstream of the next exon-exon junction ^9–11^. The EJC is bridged by UPF3A/B and UPF2 to UPF1, the central NMD factor ^12^. Subsequently, the unstructured N- and C-terminal tails of UPF1 are phosphorylated at [S/T]Q motifs by the SMG1:SMG8:SMG9 complex, which consists of the kinase SMG1 and its regulators SMG8 and SMG9^13–17^. UPF1 harbours 28 [S/T]Q motifs, of which 19 are evolutionarily conserved ^18^. While no single phosphorylation site appears to be indispensable for NMD, the synergistic effect of phosphorylation at multiple sites contributes to the degradation process, with each site having a varying degree of importance ^18^. The heterodimer of SMG5:SMG7 selectively binds to phosphorylated UPF1, whereas the endonuclease SMG6 interacts both in a phosphorylation-dependent and -independent manner with UPF1^18–22^. Nevertheless, SMG5:SMG7 are crucial for the activation of SMG6, which results in mRNA cleavage in the vicinity of the PTC^7,20,23,24^.

In metazoans, the conserved SMG1:SMG8:SMG9 complex plays an important role in ensuring the precise execution of the NMD processes. SMG1 belongs to the phosphatidylinositol-3-kinase-related kinase (PIKK) family, and phosphorylates serine or threonine residues ^17^. It prefers a glutamine residue at position +1 and leucine residue at -1 position for efficient phosphorylation ^14,17,25^. Structurally, SMG1 comprises of a catalytically active C-terminal head and an N-terminal arm including the N-HEAT domain forming an alpha-solenoid ^26^. The so-called insertion domain of SMG1 functions as PIKK-regulatory domain (PRD) and its removal leads to hyperactivation of the kinase ^27,28^. Functionally, the SMG1 insertion domain inhibits substrate binding and blocks the access to the active site ^29^.

SMG8 is composed of an N-terminal G-like domain followed by a C-terminal kinase inhibitory domain (KID), while SMG9 features a C-terminal G-domain. SMG8 and SMG9 form an unusual heterodimer with SMG9’s G-domain and SMG8’s G-like domain interaction mirroring that of dimeric GTPases^30^. On the side opposite to SMG8, the G-domain of SMG9 interacts with SMG1 via the N-HEAT domain and the alpha-solenoid ^13,25,26,30^. SMG8 interacts with SMG1 only via the alpha-solenoid, enabling SMG9 to control the activity of SMG1 indirectly via integration of SMG8 into the complex ^13,25,26,30^. Previous studies have shown that the removal of SMG8 or the deletion of its KID resulted in increased SMG1 activity, suggesting a regulatory role of SMG8 in inhibiting SMG1 via its KID^13,26–29,31^. Mechanistically, the KID stabilizes SMG1 in its autoinhibited state, offering insight into how SMG8 regulates SMG1 activity at a molecular level ^29^.

The existing knowledge about SMG8 and SMG9 is largely derived from analyzing recombinant proteins *in vitro*, leaving a gap in understanding their contributions to NMD regulation within intact cells. In an endeavour to bridge this knowledge gap, our initial hypothesis suggested that deleting the KID of endogenous SMG8 would enhance SMG1 activity, resulting in UPF1 hyperphosphorylation. However, we detected no changes in steady-state phosphorylation, which prompted us to generate SMG8 and SMG9 knock-out (KO) cells via CRISPR-Cas9, resulting in mild NMD impairment and no altered UPF1 phosphorylation. Treatment of SMG8- and SMG9-KO cells with SMG1 inhibitor (SMG1i) resulted in severe NMD impairment, but no altered UPF1 phosphorylation compared to equally treated WT cell. We analyzed the transcriptome-wide effects of SMG1i via RNA-Seq and detected concentration and KO-dependent upregulation of NMD-annotated transcripts. Analysis of the interactome of immunoprecipitated endogenous UPF1 in the absence or presence of SMG1i suggests that SMG8- and SMG9-KOs and SMG1i treatment stall inactive complexes at different stages along the assembly of the NMD machinery. Collectively, these results provide an extensive characterization of the SMG1:SMG8:SMG9 complex, which serves to maintain robustness during the first authentication step of human NMD.

## Results

### The KID of SMG8 is dispensable for NMD

A pivotal step during the initiation of NMD is the phosphorylation of PTC-proximal UPF1 molecules by the SMG1:8:9 complex, marking aberrant transcripts for degradation (Figure 1A). Subsequent binding of SMG5:SMG7 to phosphorylated UPF1 enables the endonucleolytic decay of the target mRNA by SMG6. The exact modalities of the recruitment of the SMG1:8:9 complex to UPF1 and the activation of this complex are still not fully understood. However, previous studies reported that SMG8 inhibits SMG1 *in vitro* through its C-terminal kinase inhibitory domain (KID), suggesting a potential regulatory role in UPF1 phosphorylation ^13,26–29,31^. Based on these previous findings, we hypothesized that the deletion of the KID would lead to increased UPF1 phosphorylation and altered NMD activity in cells. To test this hypothesis, we generated HCT116 cells in which a cassette containing a Myc tag, a Puromycin resistance marker, and poly(A) signal was inserted into the second exon of the endogenous SMG8 locus using the CRISPaint method ^32^ (Figure S1A, Table S1). This genomic modification results in the deletion of the SMG8 KID, which was confirmed by Western blot analysis (Figure 1B). However, the UPF1 phosphorylation level as detected by a phospho-UPF1 specific antibody (serine 1116; short loop isoform, Uniprot ID Q92900-2) was not or only slightly changed in these cells (Figure 1C).

**Figure 1.**
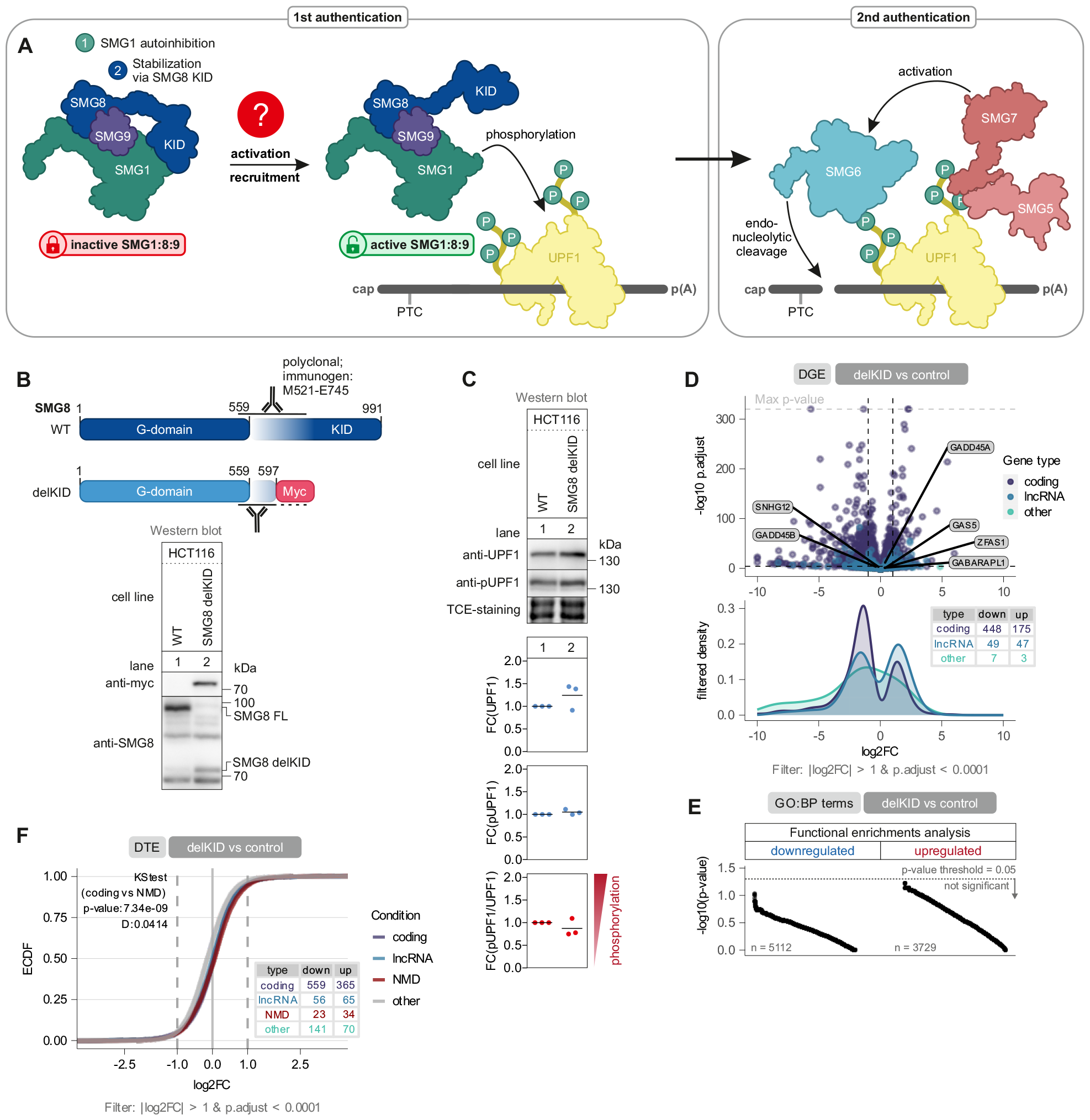
NMD activity is unaffected by the deletion the SMG8 KID. (**A**) Schematic representation of the central steps of nonsense-mediated mRNA decay (NMD). The SMG1 kinase is regulated by SMG8 and SMG9. The SMG1:SMG8:SMG9 complex is recruited to mRNA-bound UPF1 and phosphorylates the N- and C-terminal tails of UPF1 allowing the heterodimer SMG5:SMG7 and the endonuclease SMG6 to bind. SMG5:SMG7 activate SMG6, resulting in the endonucleolytic cleavage of the mRNA in the vicinity of the premature termination codon (PTC) via SMG6. (**B**) Western blot analysis of cells with deleted SMG8 KID (delKID; M1-P597). Anti-Myc (AK-106) and anti-SMG8 (AK-159) antibodies were used (n=1 biologically independent sample; see Table S1 for antibody details). The region of SMG8 detected by the SMG8 antibody is schematically depicted. (**C**) Analysis of endogenous UPF1 serine 1116 (corresponding to the UPF1 short loop isoform; Uniprot ID Q92900-2) phosphorylation status in WT and SMG8 delKID cells. TCE-staining serves as a control. Quantification of total UPF1 (anti-UPF1; AK-156) or phosphorylated UPF1 (anti-pUPF1; serine 1116; AK-146) is normalized to one representative TCE-staining and is shown as data points and mean (n=3 biologically independent samples). (**D**) (top) Volcano plot showing the differential gene expression in SMG8 delKID versus control RNA-Seq data. Genes with GENCODE-annotated gene biotypes protein-coding (purple), long non-coding (lncRNA; blue) or other (green) are indicated. The log2 fold change in gene expression is plotted against the -log10 adjusted p-value (p.adjust). Individual known NMD-targeted genes are highlighted. (bottom) Density plot showing the distribution of gene biotypes with significant changes, cutoffs were |log2FC|> 1 and p.adjust < 0.0001. Numbers of significantly changed genes per biotype are indicated in the inset table. (**E**) Functional enrichments analysis via g:profiler of significantly up- or downregulated genes in SMG8 delKID cells. The -log10(p-value) of each detected GO:BP term is plotted with a p-value threshold of 0.05. (**F**) Empirical cumulative distribution function (ECDF) plot of differentially expressed transcripts in SMG8 delKID cells versus control. Expression changes for GENCODE-annotated transcript biotypes proteincoding (purple), long non-coding (lncRNA; blue), NMD-annotated (red) and other (green) are indicated. EDCF plots shows the distribution in respect to the log2 fold change (log2FC) and significantly regulated transcripts are summarized in the inset table with the cutoffs |log2FC| > 1 and p.adjust < 0.0001. Kolmogorov-Smirnov test was performed for protein-coding versus NMD-annotated transcripts, showing the p-value and test statistic D.

To assess the consequences of KID removal, we sequenced RNA isolated from these cells. RNA-Seq analysis verified the deletion of the KID at the mRNA level, with no full-length SMG8 mRNA being detected (Figure S1B, Table S2). Differential expression analysis of poly(A)+ enriched mRNA revealed that more than twice as many coding genes were downregulated compared to upregulated genes (Figure 1D). Gene ontology analysis did not reveal any significant pathways associated with the observed differential expression patterns (Figure 1E). Of note, previously identified NMD-sensitive marker mRNAs, such as GADD45B, ZFAS1 or GABARAPL1^33–35^ remained unchanged, indicating that NMD activity is not globally altered. To test the NMD-status of the delKID cells in more detail, we turned to transcript isoform-specific analyses. Differential transcript expression analysis showed only minor changes in NMD-annotated target mRNAs, further substantiating that the KID deletion did not significantly impact NMD efficiency (Figure 1F). Further analysis of differentially expressed transcripts in the SMG8 delKID clone displayed that downregulated isoforms have longer transcript length including longer 5’ UTR, CDS and 3’ UTR length (Figure S1C).

In conclusion, our findings demonstrate that the deletion of the SMG8 KID does not result in significant changes in UPF1 phosphorylation or NMD activity. This implies that the inhibition of SMG1 by SMG8 does not affect phosphorylation levels of UPF1 in cells, raising questions about the relevance of this regulation *in vivo*.

### Minor NMD inhibition in SMG8 or SMG9 knock-out cell lines

Since no effect on NMD or UPF1 phosphorylation was detected when deleting the SMG8 KID, we wanted to investigate how SMG8 and SMG9 regulate SMG1. Both, SMG8 and SMG9 are considered to play a vital role in SMG1 inhibition. Hence, we hypothesized that the complete inactivation of SMG8 or SMG9 would lead to hyperphosphorylation of UPF1, resulting in altered NMD activity (Figure 2A). We generated SMG8- and SMG9-KO HCT116 cells, respectively, and SMG8-KO HEK293 cells via the CRISPaint system. KOs were verified by Western blotting and Sanger sequencing (Figure 2B, Figure S2A). SMG9 has been shown to integrate SMG8 into the SMG1:SMG8:SMG9 complex, but the precise role of SMG8 remained unclear. To test if SMG8 is involved in the interaction of SMG1 and SMG9, immunoprecipitation of FLAG-tagged SMG9 was performed (Figure S2B). Decreased SMG1-SMG9 binding in SMG8-KO cells compared to WT cells was observed indicating SMG8 contributes to the stability of the SMG1:SMG8:SMG9 complex.

**Figure 2.**
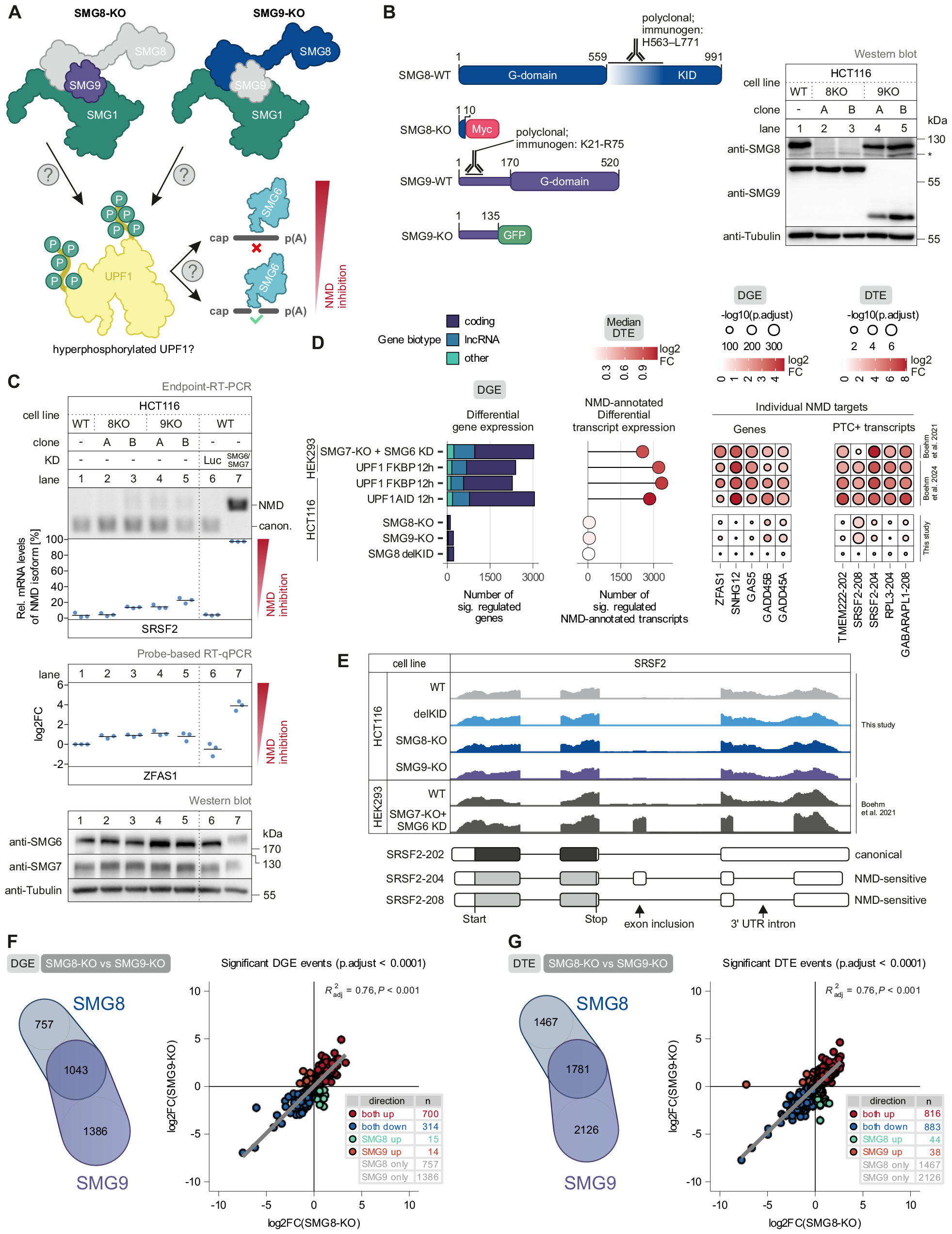
SMG8- and SMG9-KO cells show lower NMD activity. (**A**) Schematic overview of the SMG1:SMG8:SMG9 complex without SMG8 or SMG9. The lack of SMG8 or SMG9 might influence UPF1 phosphorylation resulting in altered NMD activity. (**B**) Western blot analysis of SMG8- or SMG9-KO cells using anti-SMG8 (AK-169) and anti-SMG9 (AK-170) antibodies. Tubulin (AK-084) serves as control (n=1 biologically independent sample; see Table S1 for antibody details). Asterisk indicates non-specific bands. Domain structure of full-length SMG8 and SMG9 protein and truncated proteins of SMG8- and SMG9-KO cells are shown. (**C**) End-point RT-PCR detection of SRSF2 transcript isoforms (top) and quantitative probe-based RT-PCR (bottom) of ZFAS1 in WT, SMG8-KO or SMG9-KO cells with or without indicated knock-downs (KD). The detected SRSF2 isoforms are indicated on the right (NMD = NMD-inducing isoform; canon. = canonical isoform). Relative mRNA levels of SRSF2 isoforms were quantified from bands of agarose gels (n=3 biologically independent samples). The ratio of ZFAS1 to the TBP reference was calculated; data points and means from the qPCRs are plotted as log2 fold change (log2FC) (n=3 biologically independent samples). Western blot analysis of SMG6 and SMG7 KD efficiency is shown with the anti-SMG6 (AK-135) and anti-SMG7 antibody (AK-136). TCE-staining serves as a control (n=1 biologically independent sample). (**D**) Comparison of SMG8-KO, SMG9-KO and SMG8-delKID RNA-Seq data with SMG7-KO + SMG6-KD (clone 34) ^7^ or three UPF1 degron conditions ^36^ regarding the number of significantly regulated genes (p.adjust < 0.0001 & |log2FC| > 1) stratified by GENCODE biotype (left), the number and median log2FC of significantly regulated GENCODE NMD-annotated transcripts (middle), as well as expression changes of individual NMD target genes and transcripts (right). (**E**) Read coverage of SRSF2 from SMG8-KO, SMG9-KO and SMG8-delKID and published SMG7-KO + SMG6-KD (clone 34) ^7^ RNA-Seq data is shown as Integrative Genomics Viewer (IGV) snapshots. The canonical and NMD-sensitive isoforms are schematically indicated below. (**F**,**G**) Overlaps between differentially regulated genes (F) or transcripts (G) of SMG8- and SMG9-KO cells. Scatter plots show the change in gene (F) or transcript (G) expression of SMG8-KO cells against the change in SMG9-KO cells (p.adjust < 0.0001). Linear regression with p-value (P) and adjusted coefficient of determination is shown.

To test how the depletion of SMG8 and SMG9 influences NMD activity, we analyzed the well-known NMD target SRSF2 and the two NMD-sensitive lncRNAs GAS5 and ZFAS1^34,37^. In HCT116 cells, all three NMD targets exhibited low accumulation compared to co-depletion of the NMD factors SMG6 and SMG7 that completely abolish NMD activity (Figure 2C, Figure S2C). HEK293 cells depleted of SMG8 showed similar effects for SRSF2, however, GAS5 and ZFAS1 showed stronger NMD inhibition similar to SMG7-KO cells (Figure S2D). Hence, SMG8- and SMG9-KO cells have a weak to moderate NMD inhibition. RNA-Seq of the SMG8-depleted HCT116 cells verified the complete loss of the SMG8 mRNA (Figure S2E, Table S2). The SMG9-KO cells expressed almost normal levels of the expected shortened transcript as well as low levels of alternatively spliced SMG9 mRNA. Almost all of these exon-skipping transcripts lead to frame shifts resulting in truncated and presumable non-functional transcripts (Figure S2F). Differential gene expression (DGE) and differential transcript expression (DTE) analysis of NMD-annotated transcripts revealed weak transcriptome-wide effects in SMG8-or SMG9-KO cells compared to conditions with severe NMD inhibition (SMG7-KO+SMG6 knock-down (KD) ^7^ or AID/FKBP-UPF1^36^) (Figure 2D-E). Comparison of significant DGE and DTE events revealed 1043 altered genes and 1781 altered transcripts in SMG8- and SMG9-depleted cells (Figure 2F-G). However, comparison of DGE of SMG8-KO and SMG8 delKID cells resulted into no correlation, emphasizing that the KID is dispensable for NMD (Figure S2G).

The majority of the differentially regulated genes and transcripts in both KOs were concordantly up- or downregulated, which is in line with the previous observations that SMG9 functions as a bridge between SMG1 and SMG8^13,25,26,30^, enabling SMG8 to execute its regulatory function towards SMG1. To test this explanation, we generated SMG8-KO cell lines expressing stably integrated SMG8 mutants with impaired SMG9 binding (mut 9A, mut 9B and mut 9AB) ^28^. Co-immunoprecipitation assays confirmed the impaired SMG1 and SMG9 interaction of the mutants (Figure S2H). Additionally, rescue assays were performed to assess the functional implications of this impaired interaction (Figure S2I). Cells expressing the mutants showed an NMD defect that was comparable to cells lacking SMG8. This suggests that the bridging function of SMG9, which facilitates the interaction between SMG1 and SMG8, is essential for full NMD activity. Taken together, loss of SMG8 or SMG9 resulted in mild NMD impairment suggesting a modulatory rather than an essential role in regulating SMG1 activity.

### SMG8- and SMG9-KO cells show unchanged steady-state UPF1 phosphorylation

Based on the mild NMD effects resulting from SMG8 or SMG9 depletion, we asked whether this effect is caused by altered levels of UPF1 phosphorylation. To answer this question, we performed Western blot analysis of whole cell lysates of SMG8-or SMG9-deficient cells. The result showed no clear increase or decrease in phosphorylation of UPF1 serine 1116 (Figure 3A-B). To provide further insight into the global phosphorylation status of UPF1, we established HCT116 cells with endogenously N-terminally FLAG-tagged UPF1 via the PITCh system (Figure 3C) ^38^. This allows immunoprecipitation of endogenous UPF1 followed by the application of an antibody, which detects phosphorylated SQ or TQ motifs with a preference for LSQ or LTQ. FLAG immunoprecipitation followed by detection with this phosphorylation antibody revealed no distinct upregulation of phosphorylation (Figure 3D-E). Therefore, these data indicate that SMG8 and SMG9 do not fundamentally influence UPF1 phosphorylation.

**Figure 3.**
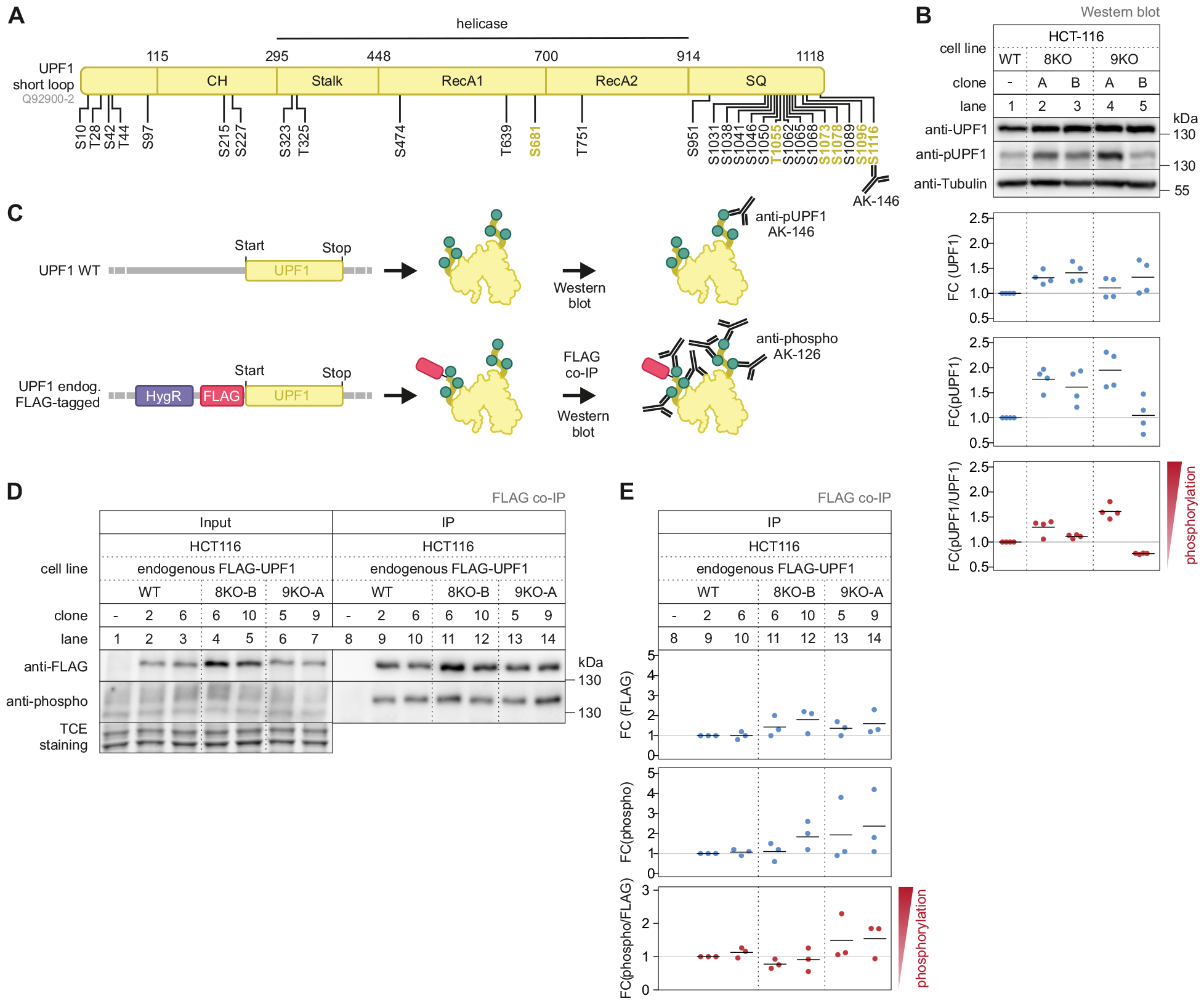
UPF1 phosphorylation is not changed in SMG8- and SMG9-KO cells. **(A)**Schematic representation of the UPF1 domain structure (short loop isoform; Uniprot ID Q92900-2). Positions of [S/T]Q motifs are indicated with black lines and black font. L[S/T]Q motifs are shown with black lines and yellow font. The epitope recognized by the pUPF1 antibody (serine 1116; AK-146) is indicated. **(B)**Analysis of endogenous UPF1 serine 1116 (corresponding to the UPF1 short loop isoform; Uniprot ID Q92900 2) phosphorylation status of whole cell lysate of WT, SMG8-KO and SMG9-KO cells. Tubulin (AK-084) serves as a loading control. Quantification of total UPF1 (anti-UPF1; AK-156) or phosphorylated UPF1 (anti-pUPF1; serine 1116; AK-146) is normalized to one representative Tubulin blot and is shown as data points and mean (n=4 biologically independent samples; see Table S1 for antibody details). **(C)** Schematic overview of the phosphorylation assay via endogenously FLAG-tagged cells generated with the CRIS-PITCh system ^38^. After FLAG co-IP, western blot analysis was performed with an serine/threonine phosphorylation-specific antibody (anti-phospho; AK-126), which preferentially binds L[S/T]Q motifs. **(D)** Western blot after FLAG co-immunoprecipitation (IP) of untagged (control) or endogenously FLAG-tagged UPF1 in WT, SMG8-KO or SMG9-KO cells. Anti-FLAG (AK-115) and anti-phospho (AK-126) antibodies were used. TCE-staining serves as a control (n=3 biologically independent samples). **(E)** Quantification of (D). Total UPF1 (anti-UPF1; AK-156) or phosphorylated UPF1 (anti-phospho; AK-126) normalized to one representative TCE-staining and is shown as data points and mean (n=3 biologically independent samples).

### Hypersensitivity to SMG1 inhibition of SMG8-or SMG9-deficient cells

The previously reported regulatory role of SMG8 and SMG9 suggests that in their absence SMG1 becomes more active (Figure 4A). Although we could not clearly detect increased UPF1 phosphorylation upon loss of SMG8 or SMG9, we explored approaches to counteract this potentially increased activity. To this end, we first downregulated SMG1 expression via siRNA-mediated KD. Contrary to expectation we observed a strong accumulation of the NMD target SRSF2 and moderate accumulation of ZFAS1 and GAS5 in SMG8- and SMG9-KO cells compared to WT cells (Figure 4B, Figure S3A). To determine if this effect was caused by the absence of the SMG1 protein itself or by the lack of its kinase function, the small molecule SMG1 inhibitor (called SMG1i) was used ^39^. SMG1i functions as an ATP-competitive inhibitor and binds to the active site of SMG1, which stabilizes the autoinhibitory conformation of SMG1^29^. Treatment of WT cells with 0.5 μM SMG1i caused severe NMD inhibition in combination with UPF1 hypophosphorylation (Figure 4C, Figure S3B-C). SMG8- and SMG9- KO cells, however, showed severe NMD inhibition already upon treatment with low concentrations (0.1 μM) of SMG1i, indicating an increased sensitivity to SMG1 inhibition in the absence of SMG8 or SMG9 (Figure 4D, Figure S3D). UFP1 phosphorylation levels were similar in WT and KO cells upon 0.1 μM SMG1i indicating that the severe NMD impairment cannot be merely explained by altered global UPF1 phosphorylation status (Figure 4E; compare lane 13,16,19). In contrast, higher concentration of SMG1i (1 μM) led to reduced phosphorylation of serine 1116 in SMG8- and SMG9- KO cells compared to WT cells, indicating (1) that SMG8 and SMG9 do influence UPF1 phosphorylation under challenging conditions and (2) that not all UPF1 phosphorylation sites have the same phosphorylation status. Taken together, these experiments demonstrate that SMG8 and SMG9 contribute to the robustness of NMD execution and that the mere UPF1 phosphorylation status does not positively correlate with NMD activity.

**Figure 4.**
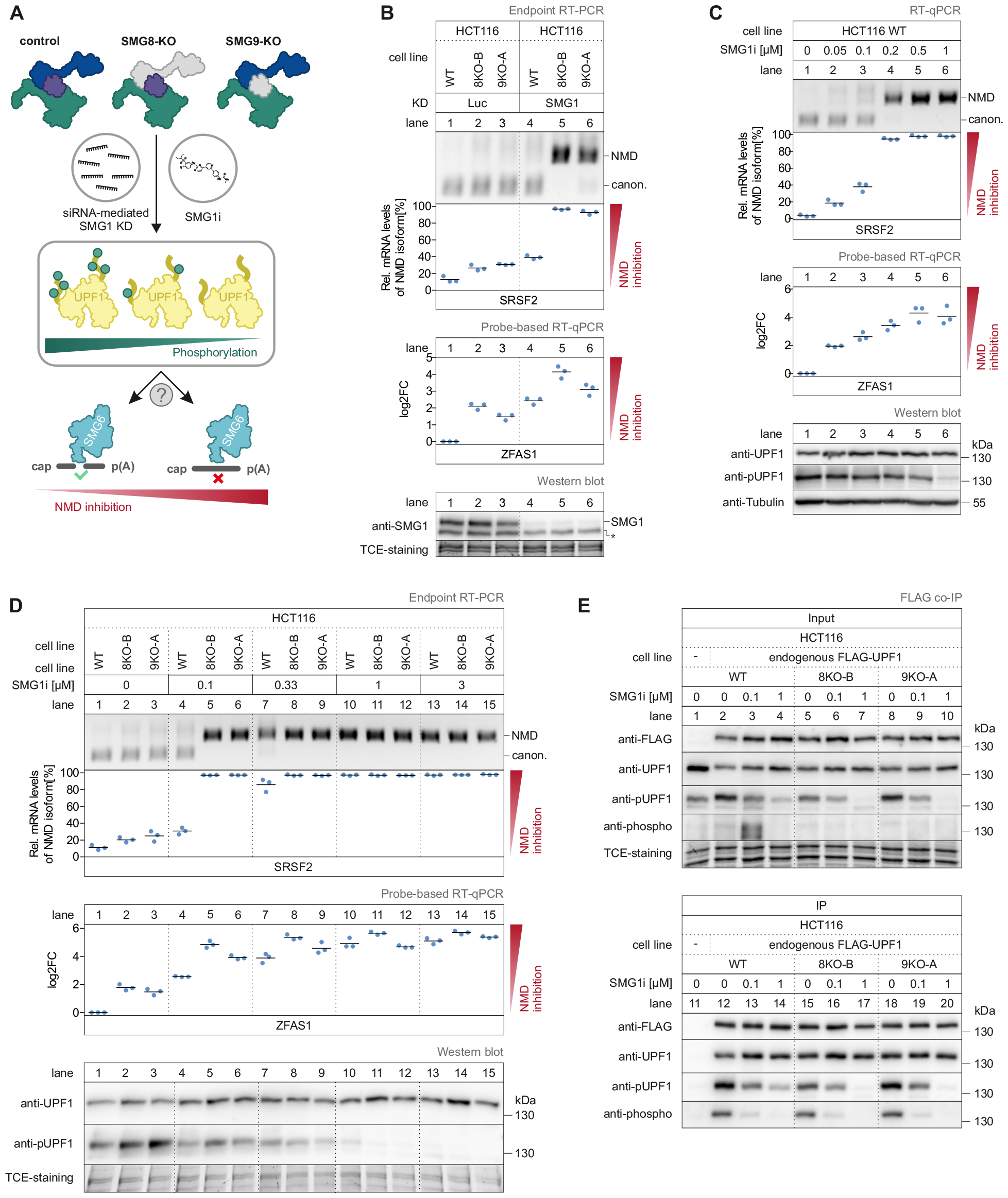
SMG8- and SMG9-KO cells are hypersensitive to SMG1i treatment. (**A**) Schematic representation of SMG1 inactivation via siRNA-mediated knock-down (KD) or treatment with the SMG1 inhibitor SMG1i and subsequent expected changes in UPF1 phosphorylation and NMD activity. (**B**) End-point RT-PCR detection of SRSF2 transcript isoforms (top) and quantitative probe-based RT-PCR (bottom) of ZFAS1 in WT, SMG8-KO or SMG9-KO cells with Luc or SMG1 KD. The detected SRSF2 isoforms are indicated on the right (NMD = NMD-inducing isoform; canon. = canonical isoform). Relative mRNA levels of SRSF2 isoforms were quantified from bands of agarose gels (n=3 biologically independent samples). The ratio of ZFAS1 to the B2M reference was calculated; data points and means from the qPCRs are plotted as log2 fold change (log2FC) (n=3 biologically independent samples). Western blot analysis of SMG1 KD efficiency is determined with the anti-SMG1 antibody (AK-088) and TCE-staining serves as a control (n=1 biologically independent sample; see Table S1 for antibody details). (**C, D**) End-point RT-PCR detection of SRSF2 transcript isoforms (top) and quantitative probe-based RT-PCR (bottom) of ZFAS1 in WT cells (C), SMG8-KO and SMG9-KO cells (D) with treatment of different SMG1i concentrations for 24 h. The detected SRSF2 isoforms are indicated on the right (NMD = NMD-inducing isoform; canon. = canonical isoform). Relative mRNA levels of SRSF2 isoforms were quantified from bands of agarose gels (n=3 biologically independent samples). The ratio of ZFAS1 to the B2M reference was calculated; data points and means from the qPCRs are plotted as log2 fold change (log2FC) (n=3 biologically independent samples). Analysis of endogenous UPF1 serine 1116 (corresponding to the UPF1 short loop isoform; Uniprot ID Q92900-2) phosphorylation status was determined with anti-UPF1 (AK-128) and anti-pUPF1 (serine 1116; AK-146). TCE-staining serves as a control (n=1 biologically independent sample). (**E**) Western blot after FLAG co-immunoprecipitation (IP) of untagged (control) or endogenously FLAG-tagged UPF1 in WT, SMG8- KO or SMG9-KO cells treated with SMG1i for 24 h. Anti-FLAG (AK-103), anti-pUPF1 (serine 1116; AK-146) and anti-phospho (binds phosphorylated serine/threonine; AK-126) were used. TCE-staining serves as a control (n=3 biologically independent samples).

### SMG1i treatment results in concentration dependent NMD inhibition

Given the strong accumulation of NMD-sensitive transcripts of SRSF2, ZFAS1 and GAS5 upon SMG1 inactivation via SMG1i, we next sought to investigate the transcriptome-wide effects caused by this treatment. Principal component analysis of RNA-Seq-derived global gene-level counts exhibited an almost linear trend with increasing concentrations of SMG1i (Figure 5A, Table S2). The delKID cells showed a similar distribution compared to WT cells, however, were shifted along principal component 2. In addition, up- and downregulated genes of cells lacking the SMG8 KID (Figure 1D) were not differentially regulated by SMG1i treatment (Figure S4A), suggesting that these were clone-specific effects.

DGE analysis of significantly regulated genes confirmed the hypersensitivity of the SMG8- and SMG9- KO cells compared to WT cells when exposed to low concentrations of SMG1i (Figure 5B). Furthermore, high concentrations of SMG1i (1 μM) resulted in strong NMD suppression, similar to the effects observed with SMG6 and SMG7 co-depletion or UPF1 depletion via degron tag. Despite nearly completely shutting down NMD, the inactivation of SMG1 did not affect the gene expression of NMD core and EJC factors (Figure S4B).

To identify the core set of genes regulated by the SMG1:SMG8:SMG9 complex, we applied criteria to identify genes that are differentially regulated in HCT116 WT, SMG8- and SMG9-KO cells after treatment with 1 μM SMG1i. The genes were clustered into different sets including core (regulated in all three conditions), shell (regulated in two conditions) or cloud (regulated in only one condition) (Figure 5C). The core set comprised of 2016 upregulated and 1003 downregulated genes. When analyzing the overlap of identified genes between different RNA-Seq data sets with deficient NMD activity (SMG567^7^, UPF3^40^ and UPF1 core ^36^), we found that 323 genes were upregulated in all conditions, classifying these genes as high confidence NMD targets (Figure 5D, Table S3). Next, we analyzed the distribution of upregulated genes and transcripts of the SMG1i core set and found similar distribution shifts in response to NMD impairment (Figure 5E, Figure S4C). In conclusion, these results demonstrate that the sensitivity of SMG8- and SMG9-depleted cells upon SMG1i treatment has transcriptome-wide effects. Furthermore, we defined high-confidence NMD-targets using the SMG1i core set.

**Figure 5.**
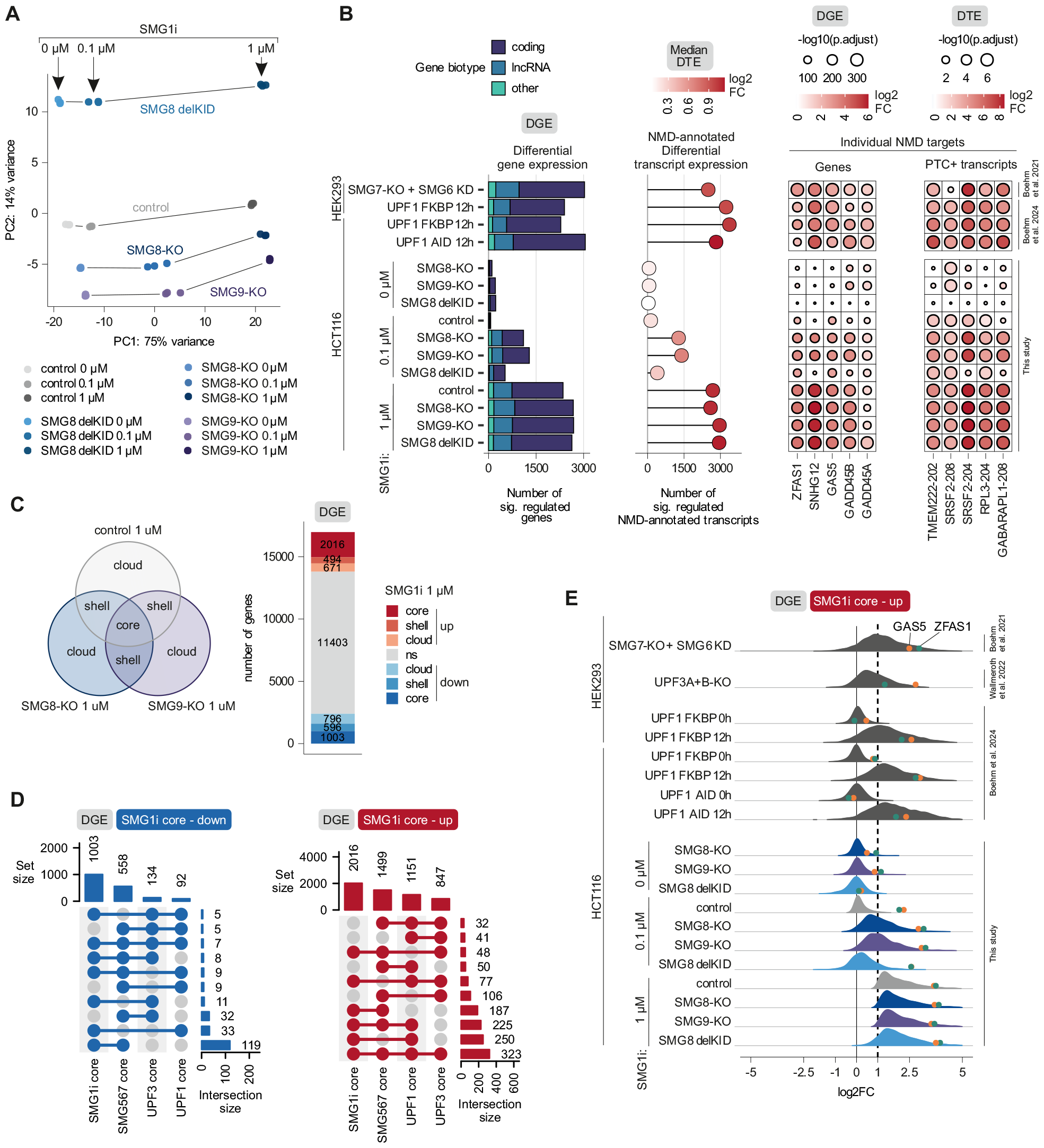
NMD is strongly inhibited transcriptome-wide after SMG1 inactivation. (**A**) Principal component analysis of gene-level counts from RNA-Seq data of WT, SMG8-KO, SMG9-KO and SMG8 delKID cells treated with different SMG1i concentrations for 24 h. Lines were added to visualize the samples from the same cell line. (**B**) RNA-Seq data of WT, SMG8-KO, SMG9-KO and SMG8 delKID cells treated with SMG1i for 24 h were compared with SMG7-KO + SMG6 knock-down (KD; clone 34) ^7^ or three UPF1 degron conditions ^36^ regarding the number of significantly regulated genes (p.adjust < 0.0001 & |log2FC| > 1) stratified by GENCODE biotype (left), the number and median log2FC of significantly regulated GENCODE NMD-annotated transcripts (middle), as well as expression changes of individual NMD target genes and transcripts (right). (**C**) Differentially expressed genes of WT, SMG8-KO and SMG9-KO cells treated with 1 μM SMG1i (significance cutoffs p.adjust < 0.0001 & |log2FC| > 1) were clustered into different sets (depicted on the left) including core (regulated in all three conditions), shell (regulated in two conditions) or cloud (regulated in only one condition). Absolute numbers of genes per set are shown on the right. (**D**) UpSet plots of the overlap of significantly up- or downregulated transcripts (p.adjust < 0.0001 & |log2FC| > 1) between SMG1i core, SMG567 core ^7^, UPF3 core ^40^ and UPF1 core ^36^. (**E**) Distribution of expression changes of the upregulated SMG1i core genes in RNA-Seq data obtained in this study and compared to SMG7-KO + SMG6-KD (clone 34) ^7^, UPF3A/B-KO ^40^ and three UPF1 degron conditions ^36^. Previously used reporter genes GAS5 (orange) and ZFAS1 (green) are indicated as circles.

### Catalytically inactive SMG1:SMG8:SMG9 complexes accumulate in association with UPF1

The inactivation of SMG1 led to UPF1 hypophosphorylation, which is expected to affect the interactome of UPF1. This prompted us to conduct immunoprecipitation of endogenously FLAG-tagged UPF1, followed by label-free mass spectrometry analysis. In untreated (no SMG1i) HCT116 WT cells almost all NMD and EJC factors as well as both Staufen proteins (STAU1, STAU2) were enriched (Figure 6A-B, Figure S5A, Table S4). However, other previously identified UPF1 interaction partners (like PNRC2) were not detected ^41^(Figure S5B). Treatment with SMG1i resulted in an increased presence of SMG1, SMG8 and SMG9 in the UPF1 interactome, suggesting that UPF1 has to undergo phosphorylation by SMG1 to facilitate the efficient recycling of the SMG1:SMG8:SMG9 complex (Figure 6B). Furthermore, decreased levels of SMG5 and SMG7 were detected at high concentrations of SMG1i, which is well in line with their phosphorylation-dependent mode of interaction. Although NMD was completely abolished under this condition, the endonuclease SMG6 was enriched, underlining (1) the ability of SMG6 to bind to UPF1 independently of its phosphorylation and (2) the requirement of SMG5:SMG7 to activate SMG6. In addition, all core EJC factors were enriched following SMG1i treatment, indicating that the NMD machinery dissociates slower from its target mRNA.

**Figure 6.**
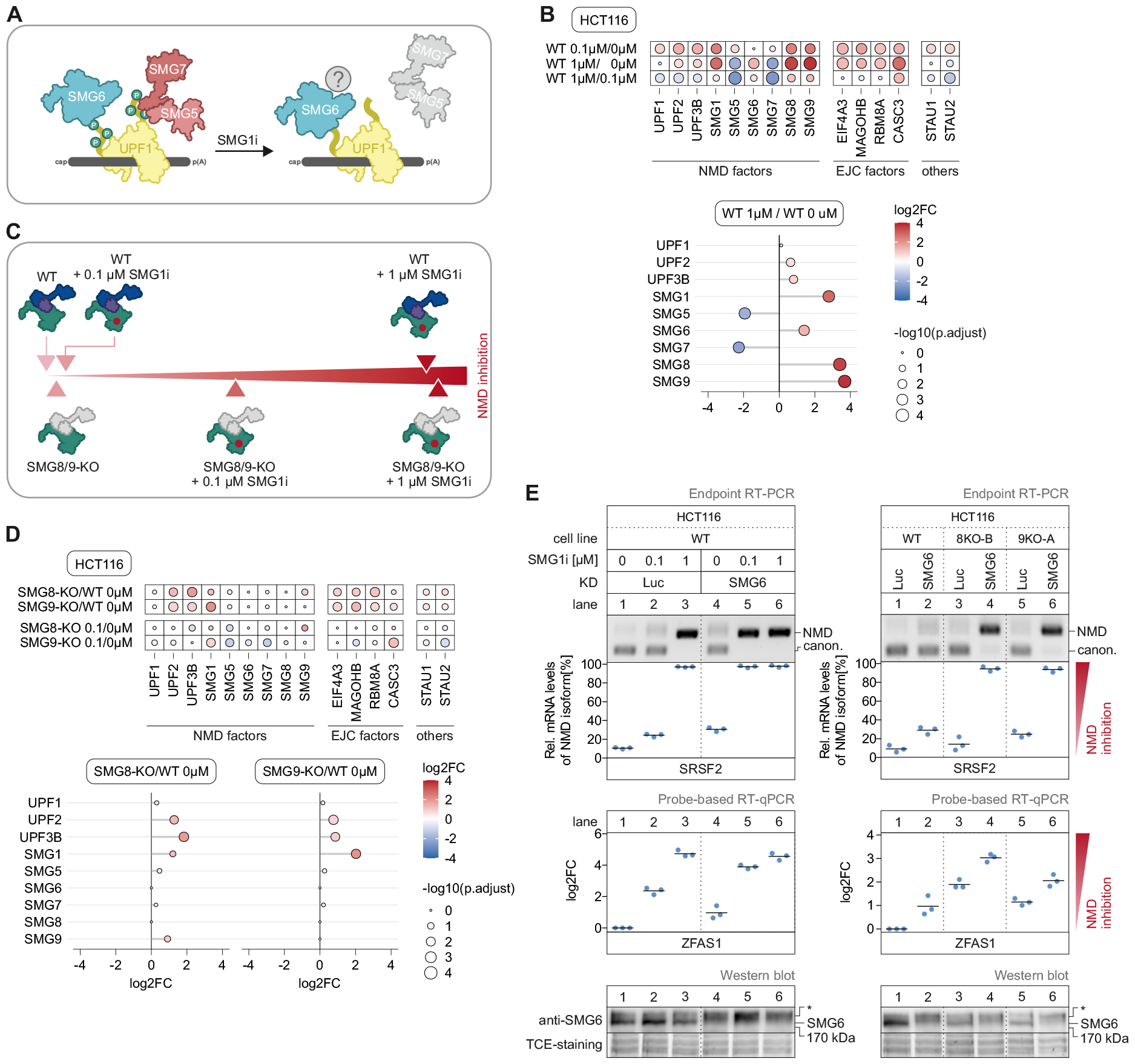
SMG1i-treated cells show an increased association of UPF1 with the SMG1:SMG8:SMG9 complex. (**A**) Schematic overview of UPF1 binding partners after SMG1i treatment. (**B**) Heatmap of mass spectrometry-based analysis of FLAG co-immunoprecipitated (IP), untagged (control) or endogenously FLAG-tagged UPF1 in WT cells. Cells were treated with indicated concentrations of SMG1i for 24 h. Colored points indicate the log2 fold change (log2FC) and point size corresponds to the adjusted p-value (adj. p-value; from Students t test; n = 4 biologically independent samples). (**C**) Schematic overview of the NMD inhibition of WT, SMG8-KO or SMG9-KO cells upon SMG1i treatment. (**D**) Heatmap of mass spectrometry-based analysis of FLAG co-immunoprecipitated (IP), untagged (control) or endogenously FLAG-tagged UPF1 in SMG8-KO and SMG9-KO cells. Cells were treated with the indicated concentrations of SMG1i for 24 h. Colored points indicate the log2 fold change (log2FC) and point size corresponds to the adjusted p-value (adj. p-value; from Students t test; n = 4 biologically independent samples). (**E**) End-point RT-PCR detection of SRSF2 transcript isoforms (top) and quantitative probe-based RT-PCR (bottom) of ZFAS1 in WT, SMG8-KO and SMG9-KO cells with Luc knock-down (KD; control) or SMG6-KD. WT cells were treated in addition with SMG1i for 24 h. The detected SRSF2 isoforms are indicated on the right (NMD = NMD-inducing isoform; canon. = canonical isoform). Relative mRNA levels of SRSF2 isoforms were quantified from bands of agarose gels (n=3 biologically independent samples). The ratio of ZFAS1 to the B2M reference was calculated; data points and means from the qPCRs are plotted as log2 fold change (log2FC) (n=3 biologically independent samples). Western blot analysis of SMG6-KD efficiency is shown with the anti-SMG6 antibody (AK-135). TCE-staining serves as a control (n=1 biologically independent sample; see Table S1 for antibody details).

SMG8- and SMG9-KOs resulted in an increased binding of SMG1 to UPF1, indicating that both factors negatively influence S MG1-UPF1 binding (Figure 6C-D). However, binding of UPF1 to SMG1 was more pronounced in SMG9-KO cells compared to SMG8-KO cells. One possible explanation for this could be that in SMG9-KO cells also SMG8 is unable to bind to SMG1 due to the absence of SMG9’s bridging function. In addition to increased SMG1 binding, depletion of SMG8 and SMG9 resulted into accumulation of UPF2, UPF3B and EJC factors to UPF1, suggesting that both SMG8 and SMG9 contribute directly or indirectly to the dissociation of the NMD machinery (Figure 6D). Notably, only minor changes were detected in the UPF1 interactome of SMG1i-treated SMG8- and SMG9-KO cells compared to WT cells. These included reduced interactions of UPF1 with SMG5 and SMG7 (Figure S5A) despite no changes in UPF1 phosphorylation status (Figure 4E). Taken together, the catalytically inhibited SMG1:SMG8:SMG9 complex exhibits increased binding affinity to UPF1 suggesting that its phosphorylation is needed for the dissociation of the complex. In addition, reduced NMD leads to the stalling of the NMD complex.

Our previous results indicate that SMG8 and SMG9 function as modulators of SMG1 activity. While they are not strictly essential for NMD, their absence increases the sensitivity of NMD to further perturbations. We aimed to determine whether this effect is specific to the SMG1:SMG8:SMG9 complex or if it also influences later stages of NMD. In this case, cells with a perturbed SMG1:SMG8:SMG9 complex should exhibit sensitivity to the depletion of the NMD-executing factor SMG6. We explored this hypothesis through two different approaches, either with SMG1i treatment or using SMG8- or SMG9-KO cells. In WT cells combining 0.1 μM SMG1i treatment with siRNA-mediated KD of SMG6 resulted in substantial NMD inhibition (Figure 6E, Figure S5C). Similarly, SMG6-KD in SMG8- and SMG9- KO cells resulted in full NMD inhibition when analysed using SRSF2 as NMD substrate. However, only partial NMD inhibition was observed for ZFAS1 and GAS5 (Figure 6E, Figure S5C, lower part), underlining the different sensitivities of NMD targets. These results demonstrate that perturbations of the NMD machinery at early steps also affect the robustness of later stages. Thereby, they support the two-factor authentication model, which suggests that at least two authentication steps are required to license the execution of NMD.

## Discussion

Before an NMD substrate undergoes degradation by the NMD machinery, several conditions must be met. Of these requirements, the phosphorylation of UPF1 is considered a key step, previously regarded as the point of no return for mRNA degradation ^7,42–44^. However, the process of UPF1 phosphorylation via SMG1 and its regulators SMG8 and SMG9 remains incompletely understood. In this work, we systematically investigated this knowledge gap using KO cell lines of SMG8 and SMG9 as well as pharmacological inhibition of SMG1.

Previous studies on the SMG1:SMG8:SMG9 complex demonstrated that a SMG1 complex lacking SMG8 exhibits increased phosphorylation activity. This suggested that SMG8 plays an inhibitory role, which is mediated via its C-terminal KID^13,27–29,31^. We initially examined this inhibitory role using a cell line where the KID of the endogenous SMG8 was deleted (Figure 1). Since this cell line showed no change in UPF1 phosphorylation, we additionally generated SMG8- and SMG9-KO cells (Figure 2). However, even within these cells, we failed to observe the anticipated and previously reported increase in UPF1 phosphorylation (Figure 1, Figure 3). One potential reason for this discrepancy could be that the previous studies primarily relied on purified SMG1:SMG8:SMG9 complexes in *in vitro* assays. Although these assays have many advantages, they fail to capture the inherent complexity and dynamics of a living organism. The regulation of NMD involves diverse proteins, including phosphatases, which contribute to the physiological phosphorylation state of UPF1, but are absent in *in vitro* assays. It is conceivable that the depletion of SMG8 and SMG9 is accompanied by a transient increase in UPF1 phosphorylation. However, if this is counterbalanced by increased dephosphorylation, the effect of SMG8 and SMG9 depletion on UPF1 phosphorylation will be undetectable in cells. In addition to phosphatases, other factors such as UPF2 contribute to NMD regulation. UPF2 interacts with SMG1 and UPF1, but its exact role during SMG1 regulation is not clear ^42,45^. It was shown that UPF2 contributes to SMG1 activation, but also destabilizes the SMG1:SMG8:SMG9:UPF1 complex, adding another level of regulation to UPF1 phosphorylation ^16,27,46^. UPF2 is absent from purified SMG1, which could result in increased UPF1 phosphorylation in the absence of SMG8 *in vitro*. However, in *in vivo* assays the presence of UPF2 could buffer the effect of SMG8 depletion by destabilizing the binding of more active SMG1 to UPF1. One way to investigate these possibilities would be to purify the SMG1 complex from different cells (WT, SMG8-KO, SMG9-KO) and measure its kinase activity *in vitro*.

In contrast to SMG8 and SMG9, SMG1 is an essential gene in cultivated human cells. Instead of a SMG1- KO, we used the pharmacological inhibitor SMG1i, which leads to SMG1 inhibition, UPF1 hypophosphorylation and NMD inhibition. SMG1i was originally developed to enable the expression of truncated, but partially active proteins from nonsense-mutated mRNAs, which can prevent or milden symptoms of patients as shown for CFTR mRNA in cystic fibrosis ^47^. SMG1i and other NMD-inhibiting drugs have also promising applications in oncology, where NMD eliminates cancer-specific, PTC-containing transcripts and thereby prevents the production of aberrant tumour-specific neoantigens ^48^. Consequently, the inhibition of NMD factors can suppress tumour growth *in vivo* ^49,50^. Similarly, the impairment of SMG1 via the inhibitor KVS0001 increased the expression of cancer neoantigens, which can induce a T cell-dependent immune response ^51^. The physiological significance of the SMG1:SMG8:SMG9 complex extends beyond these instances. Knock-out of SMG1 or SMG9 in mice results in embryonic lethality, highlighting their crucial roles in development ^52,53^. Additionally, homozygous loss-of-function mutations of the SMG8 or SMG9 gene in patients are associated with various disorders, including severe developmental delay and malformations in the heart and eyes^53,54^. These examples emphasize not only the importance to understand NMD in general to treat patients with NMD-related diseases, but also to understand the impact and consequences of NMD-specific drugs in living organisms. When applying low amounts of SMG1i to SMG8- or SMG9-deficient cells, we found that they were hypersensitive to SMG1 inhibition. The identical concentration of SMG1i had minimal impact on WT cells. We suggest that during minor disturbances of NMD (e.g. SMG8-KO, SMG9-KO or small amounts of SMG1i), NMD has compensatory mechanism to preserve the physiological state of the cell. However, further disruptions of the NMD machinery exceed the limits of these mechanisms leading to synergistic NMD inhibition, as seen as hypersensitivity of SMG8- and SMG9-KO cells upon SMG1i treatment. This suggests that patients harbouring mutations in NMD factor-encoding genes, such as SMG8 and SMG9, may experience significant NMD dysfunction, potentially leading to severe side effects upon SMG1i treatment.

Our transcriptome-wide analysis demonstrates that neither SMG8 nor SMG9 are absolutely essential for NMD (Figure 5). However, SMG8-KO cells exhibited stronger NMD inhibition than SMG8 delKID cells indicating that the KID is not the only SMG8 domain contributing to NMD regulation. Although mass spectrometry analysis showed that neither SMG8 nor SMG9 are necessary to recruit SMG1 to UPF1 (Figure 6), both factors seem to serve an auxiliary function in contributing to the robustness of NMD and in their absence some NMD substrates are not efficiently degraded. The question arises as to how to explain this observation. Could this be the consequence of an exceptionally high NMD efficiency in vertebrates, which is based on a particularly intricate NMD machinery? As component of this machinery SMG8 binds via its KID to the C-terminal insertion domain of SMG1, thereby supporting the autoinhibitory state of SMG1^29^. SMG9, on the other hand, is required for SMG8 binding to SMG1. This hypothesis finds support from *C. elegans*, where the insertion domain of SMG1 is not present and depletion of SMG8 does not influence NMD ^55,56^. This could suggest that NMD in *C. elegans* generally operates with lower efficiency. Furthermore, this indicates that SMG8 and the SMG1 insertion domain contribute to a more complex regulation of UPF1 phosphorylation in vertebrates. In addition, the phosphorylation of UPF1 may be essential for the intricate regulation of the NMD machinery (including compensatory mechanisms), since several lower eukaryotes are lacking SMG1 completely such as yeast, ciliates and fungi ^57,58^.

Another potential function of SMG8 and SMG9 could also be associated with the high efficiency of NMD in vertebrates. To restrict NMD activity to authentic NMD substrates, SMG8 and SMG9 assist SMG1 in identifying the correct UPF1 proteins and phosphorylating only those bound to transcripts containing PTCs, while avoiding phosphorylation of UPF1 proteins bound to normal transcripts. During SMG8 or SMG9 depletion, SMG1 could promiscuously phosphorylate all UPF1 proteins rather than just those associated with NMD-targets. Since our phosphorylation assay cannot differentiate between these UPF1 proteins, this could explain why the overall UPF1 phosphorylation status remains unchanged in SMG8- or SMG9-depleted cells. However, we currently lack evidence for this theory because the mRNAs which are downregulated in the delKID clone (Figure 1D) are not rescued by treatment with low concentrations of SMG1i (Figure S4B).

Yet another potential function of SMG8 and SMG9 may occur after successful phosphorylation, where they contribute to the dissociation of the SMG1:SMG8:SMG9 complex from UPF1. Consistent with this, we observed an increased interaction of UPF1 with SMG1 in SMG9-KO cells (Figure 6B). After phosphorylation, SMG8 and SMG9 could support the autoinhibitory state of SMG1 allowing SMG1 to move away from the target mRNA and to search for the next unphosphorylated UPF1 associated with an NMD target. The dissociation of SMG1 might also be needed for the disassembly of other NMD factors, such as UPF2, UPF3 and the EJC, since UPF1 showed an increased interaction with these factors in our SMG8- and SMG9-KO cells (Figure 6B). One possible explanation for this could be that UPF1 transitions into removing and recycling RNA-bound NMD factors via its helicase activity after dissociation of SMG1 and execution of endocleavage. However, during depletion of SMG8 or SMG9, the UPF1 transition might not be properly initiated. In addition, the process of UPF1 phosphorylation seems to be crucial for SMG1:SMG8:SMG9 to dissociate from UPF1, since an increased interaction of UPF1 with SMG1, SMG8 and SMG9 was observed when SMG1 activity was abolished via SMG1i (Figure 6A). Furthermore, this shows that catalytically inactive SMG1 is able to interact with UPF1, which is in line with previous finding ^29^.

While UPF1 phosphorylation is an essential step during NMD execution, we found here that the level of phosphorylation does not clearly correlate with NMD activity. WT, SMG8-KO and SMG9-KO cells treated with 0.1 μM SMG1i showed similar UPF1 phosphorylation, but severe differences in NMD activity (Figure 4D-E). The importance of UPF1 phosphorylation is based on its essential role for the binding of the heterodimer SMG5 and SMG7^18,21,22^. This is also confirmed in our mass spectrometry analysis since both factors were lost from the UPF1 interactome when cells were treated with SMG1i resulting in hypophosphorylated UPF1. In contrast, SMG6 showed increased binding to UPF1 in SMG1i conditions, which is in line with a phospho-independent binding of SMG6 to UPF1^19,20^. The increased binding of SMG6 could be the result of the complete NMD inhibition. During this NMD inactivation, UPF2, UPF3B and all EJC factors (EIF4A3, MAGOH, RBM8A, CASC3) were enriched indicating a stalled NMD machinery that is incapable of recycling its factors. Interestingly, the same effect was seen in SMG5- and SMG7-depleted cells, where proximity labelling of UPF1 revealed an enrichment of UPF2, UPF3B, SMG1, SMG6, SMG8 and SMG9^7^. Notably, a shared characteristic between this condition and the NMD inhibition via SMG1i is the lack of binding of the heterodimer SMG5:SMG7 to UPF1, either due to its depletion or the absence of UPF1 phosphorylation. As previously demonstrated, SMG5 and SMG7 are required for the activation of SMG6^7^ and successful cleavage of SMG6 might be necessary for dissociation from UPF1. Alternatively, SMG5:SMG7 are not only responsible for SMG6 activation but also for the dissociation of SMG6 from UPF1. Both scenarios would result in trapped SMG6, which, due to its low abundance, becomes unavailable to the cell, resulting in NMD inactivation.

Upon integrating all available data, a comprehensive insight into the function of SMG8 and SMG9 within NMD emerges (Figure 7). In the context of the previously proposed two-step authentication model, the phosphorylation of UPF1 by the SMG1:SMG8:SMG9 complex serves as the first authentication step. Subsequently, phosphorylated UPF1 recruits SMG5:SMG7 to activate endocleavage by SMG6, constituting the second authentication step. When SMG8 or SMG9 are absent, UPF1 phosphorylation initially appears unaffected. However, the first authentication step is less robust compared to WT cells, leading to increased sensitivity against further disruptions and resulting in an accumulation of NMD factors on the mRNA. Furthermore, complete deactivation of SMG1 disrupts UPF1 phosphorylation, preventing SMG5:SMG7 from binding to UPF1. Although SMG6 can still bind to UPF1 independently of phosphorylation, it loses its endonuclease activity. As a consequence, NMD complexes persistently associate with the substrate mRNA, impeding its degradation and clearance from the cell.

**Figure 7.**
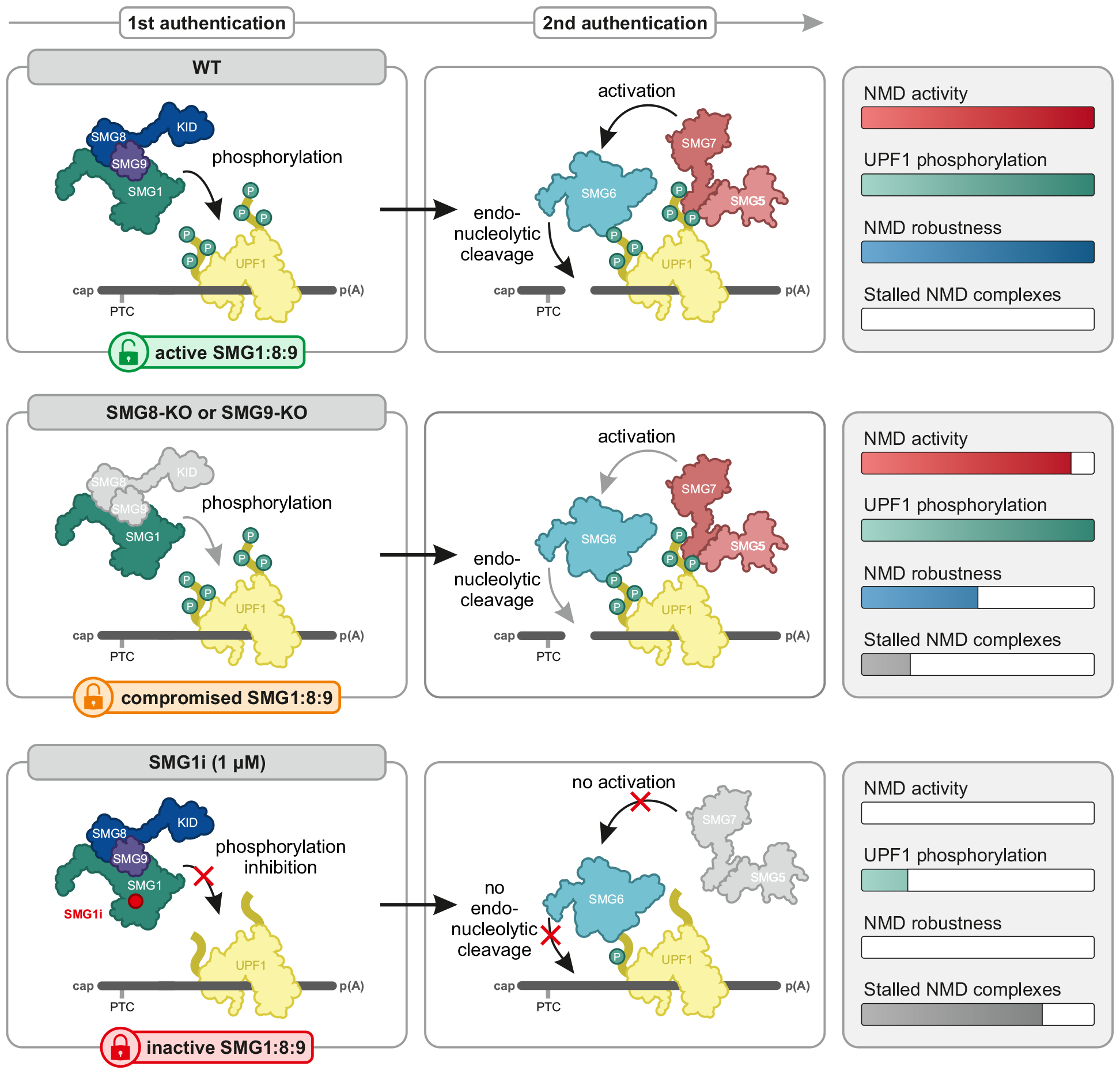
Impact of SMG1:SMG8:SMG9 complex impairment on NMD parameters. For NMD activation UPF1 has to undergo two authentication steps. (top) In wildtype cells (WT), an active SMG1:SMG8:SMG9 complex phosphorylates UPF1, which allows the SMG5:SMG7 to bind and activate SMG6. SMG6 cleaves the mRNA and the NMD complexes unbind. (middle) During the depletion of SMG8 or SMG9, SMG1 phosphorylates UPF1 and enables SMG5:SMG7 to bind and SMG6 to degrade the mRNA. However, the NMD machinery shows lower robustness against further disruptions of the NMD machinery and an accumulation of NMD factors on the mRNA. (bottom) The inactivation of SMG1 abolishes UPF1 phosphorylation. This disables SMG5:SMG7 to bind to UPF1. SMG6 can bind to UPF1 phosphorylation-independent, but remains inactive. The NMD complexes are unable to dissociate from the mRNA.

Taken together, our work reveals a previously unrecognized level of complexity in the regulation of NMD as we identify contributions of SMG8 and SMG9 to maintaining the robustness of NMD in human cells. NMD is a process with multiple layers of protection, exhibiting robust functionality even in the presence of minor disruptions. Contributing to this robustness are not only SMG8 and SMG9, but also redundant factors positioned elsewhere along the pathway, such as the paralogs UPF3A and UPF3B, or the duplicated EJC factor genes MAGOH and MAGOHB. Decreased robustness does not immediately result in dramatic consequences for NMD. Instead, it merely reduces its capacity and error tolerance, which, in turn, only manifests as phenotypically observable effects under particularly challenging conditions. This may also account for the disease symptoms associated with mutations in SMG8 and SMG9, which do not correlate with strong NMD inhibition. It is plausible that only certain tissues at certain times of development require a particularly robust NMD to develop normally, highlighting the contextual importance of NMD robustness. Hence, our findings provide not only insights into the molecular mechanisms that govern NMD regulation, but also offer potential avenues for therapeutic intervention in diseases linked to NMD dysregulation.

## Acknowledgements

We thank members of the Gehring lab for discussions and reading of the manuscript. In addition, we are thankful to Akio Yamashita for sharing the SMG8 and SMG9 antibodies. We would like to acknowledge Robert Bridges (Rosalind Franklin University of Medicine and Science) and the Cystic Fibrosis Foundation for providing the SMG1 inhibitor. We thank Veit Hornung for sharing the CRISPaint plasmids with us and Peter Kaiser, James Bradner and Behnam Nabet for providing the CRIS-PITCh plasmids. We would like to thank the CECAD Proteomics Facility for the analysis of proteome data. This work was supported by grants from the Deutsche Forschungsgemeinschaft (GE 2014/6-2) and the Center for Molecular Medicine Cologne (CMMC) (grant C 05) to N.H.G.. Additionally, this work was supported by the DFG Research Infrastructure as part of the Next Generation Sequencing Competence Network (project 423957469) and by the large instrument grant INST 216/1163-1 FUGG from the DFG (DFG Großgeräteantrag).

## Author Contributions

Conceptualization: N.H.G., S.K., V.B. Methodology: S.K., V.B., N.H.G. Software: V.B., S.K., Investigation: S.K., V.B., S.T. Resources and data curation: V.B., J.- W.L., S.K., M.F., K.B. Writing - original draft, review, and editing: S.K., N.H.G., V.B. Visualization: S.K., V.B. Supervision: N.H.G. Funding acquisition: N.H.G.

## Methods

### RESOURCE AVAILABILITY

#### Lead contact

Further information and requests for reagents should be directed to and will be fulfilled by the Lead Contact, Niels H. Gehring.

#### Materials availability

All cell lines and plasmids generated in this study are listed in Table S1 and are available upon request to the lead contact.

#### Data and code availability

- This study analyzes publicly available data, which are listed below and in the Table S1.
- RNA-Seq data generated in this study have been deposited at BioStudies/ArrayExpress and are publicly available as of the date of publication. Accession numbers are listed in Table S1.
- All original code has been deposited at GitHub (https://github.com/boehmv/2024_SMG189) and is publicly available as of the date of publication.
- The mass spectrometry proteomics data have been deposited to the ProteomeXchange Consortium via the PRIDE^59^ partner repository with the dataset identifier PXD051058.
- Any additional information required to reanalyze the data reported in this study is available from the lead contact upon request.

### EXPERIMENTAL MODEL AND SUBJECT DETAILS

#### Cell lines

HCT-116 (human, male, colorectal carcinoma, epithelial; ATCC, cat. no. CCL-247; RRID:CVCL_0291) and Flp-In-T-REx-293 (human, female, embryonic kidney, epithelial; Thermo Fisher Scientific, cat. no. R78007; RRID:CVCL_U427) were cultivated in high glucose, GlutaMAX DMEM (Gibco) supplemented with 9% fetal bovine serum (Gibco) and 1x Penicillin-Streptomycin (Gibco). The cells were cultured at 37 °C and 5% CO_2_ in a humidified incubator. The generation of knock-in/knock-out and stable cell lines is described below. All cell lines are summarized in Table S1.

### METHOD DETAILS

#### Generation of knock-out and knock-in cells using CRISPaint or CRIS-PITCh system

The SMG8 and SMG9 knock-outs/knock-ins cells were generated via the CRISPaint system ^32^. The sgRNA sequence for SMG8 delKID was 5’- CTATTGTGATATAGCACAGG-3’, for SMG8-KO 5’- AGCTTGCGAGACCTTCTAAT-3’ and for SMG9-KO 5’-TGCGCCACCCAAGGGGGAGA-3’. 2.5 x 10^5^ cells per sgRNA were seeded in 6-well plate. One day after seeding 2000 ng universal donor (pCRISPaint- myc-PuroR, Addgene plasmid # 80961; pCRISPaint- TagGFP2-PuroR, Addgene plasmid # 80970; both vectors were a gift from Veit Hornung), 1000 ng frame selector (pCAS9-mCherry-Frame +0/+1/+2, Addgene plasmid #66939/#66940/#66941; all three plasmids were a gift from Veit Hornung) and 1000 ng target selector (px330-SMG8 delKID-R57; px330-SMG8- IDT-AA; px330-SMG9-chop94) were transfected using Lipofectamine 2000 (Thermo Fisher Scientific) according to the manufacturer’s protocol. 2 days after transfection the cells were transferred to 10 cm dishes and 4-5 days after transfection cells were selected with 0.75-1.0 μg/ml Puromycin (InvivoGen). All CRISPaint plasmids are summarized in Table S1.

UPF1 was endogenously FLAG-tagged via the CRIS-PITCh v2 system ^38^. The plasmid px330- BbsI-PITCh UPF1 N is based on pX330-BbsI-PITCh (Addgene plasmid #127875; was a gift from Peter Kaiser) and encodes the UPF1 specific sgRNA (5’-CCCGTACGCCTCCACGCTCA-3’). The donor plasmid pCRIS-PITChv2-PurR-FLAG is based on pCRIS-PITChv2-dTAG-Puro (BRD4) (Addgene plasmid #91796; was a gift from James Bradner & Behnam Nabet) and contains two 40 bp-long N-terminal UPF1 microhomologies (5’-GCAGCGCGGAACCGGCCCGA GGGCCCTACCCGGAGGCACC-3’ and 5’-GAGCGT GGAGGCGTACGGGCCCAGCTCGCAGACTCTCACTT-3’) flanking a Hygromycin resistance gene, a T2A signal, the FLAG-tag, and a linker region. For transfection, 2.5 x 10^5^ cells were in seeded in 6-well plate and after 24 h 1000 ng px330-BbsI-PITCh-UPF1-N and 500 ng donor plasmid pCRIS-PITChv2-PurR-FLAG were transfected using Lipofectamine 2000 (Thermo Fisher Scientific) according to the manufacturer’s protocol. 2 days after transfection the cells were transferred to 10 cm dish and 4-5 days after transfection cells were selected with 100 μg/ml Hygromycin (InvivoGen). All CRIS-PITCh plasmids are summarized in Table S1.

For both CRISPaint and CRIS-PITCh system cells were selected for 2-3 weeks with 750 ng/ml Puromycin or 100 μg/ml Hygromycin. Cell colonies originating from a single clone were isolated in 12-well plates and genomic DNA was extracted using QuickExtract DNA Extraction Solution (Lucigen) according to manufacturer’s instruction. Correct insertion of the gene cassette was screened via genomic PCR and verified via Sanger sequencing (Eurofins Genomics). The primers for genomic PCR are listed in Table S1.

#### DNA and RNA extraction

One day prior to DNA extraction, cells were seeded in a 48-well plate. To extract DNA, 50 μl QuickExtract DNA Extraction Solution (Lucigen) was used following the manufacturer’s instructions. For RNA extraction, cells were dissolved in 1 ml in-house prepared TRI reagent ^60^ per 6-well and RNA was extracted following instructions of peqGOLD TriFast (VWR Peqlab; v0815_e). Following changes were made: Instead of 200 μl chloroform, 150 μl 1-Bromo-3-chloropropane (Sigma Aldrich) was used. RNA was resuspended in 20 μl RNase-free water.

#### Immunoblot analysis

For SDS-polyacrylamide gel electrophoresis and immunoblot analysis protein samples were harvested with RIPA buffer (50 mM Tris/HCl pH 8.0, 0.1% SDS, 150 mM NaCl, 1% IGEPAL CA-630, 0.5% deoxycholate) or samples were eluted from Anti-FLAG M2 magnetic beads (Sigma-Aldrich). For analyzation of UPF1 phosphorylation status RIPA buffer was supplemented with 1x PhosSTOP (Roche), 1x Halt Protease and Phosphatase Inhibitor Cocktail (Thermo Scientific) and 10 μg/μl RNase (Panreac AppliChem). Pierce Detergent Compatible Bradford Assay Reagent (Thermo Fisher Scientific) was used for protein quantification. All antibodies are listed in Table S1 and were used at the indicated dilutions in 50 mM Tris [pH 7.2], 150 mM NaCl with 0.2% Tween-20, and 5% skim milk powder. Amersham ECL Prime or Select Western Blotting Detection Reagent (GE Healthcare) in combination with the Fusion FX-6 Edge system (Vilber Lourmat) and Evolution-Capt Edge software (version 18.05) was used for visualization. Quantification of detected protein bands was performed in a semi-automated manner using the Image-Quant TL 1D software (version 8.1) with a rolling-ball background correction. The control condition was set to unity, quantification results are shown as data points and mean.

#### Stable cell lines and plasmids

The point and deletion mutants of SMG8 were PCR amplified using Q5 polymerase (NEB) and inserted with an N-terminal FLAG-tag via NheI and NotI restriction sites into the tetracycline-inducible pcDNA5/FRT/TO vector (Thermo Fisher Scientific). For stable integration, the Flp-In T-REx system was used: 2.5-3.0 × 10^5^ cells were seeded in 6-well plates and after 24 h 2000 ng pcDNA5 construct and 1500 ng Flp recombinase expressing plasmid pOG44 were transfected using the calcium phosphate method. 48 h after transfection, cells were transferred into 10 cm dishes and selected with 100 μg/ml Hygromycin (InvivoGen). Colonies were pooled after 15-20 days. Protein expression was induced with 1 μg/ml doxycycline. All plasmids used in this study are listed in Table S1.

#### Reverse transcription, end-point RT-PCR

1-4 μg of total RNA was used for reverse transcription in a 20 μl reaction volume with 10 μM VNN-(dT)20 primer using the GoScript Reverse Transcriptase (Promega) following the manufacturer’s instructions. For end-point PCRs, 2% of cDNA (template), 0.2 μM final concentration of sense and antisense primer (see Table S1 for sequences) and MyTaq Red Mix (Bioline) was used. After 30 PCR cycles, the PCR products were resolved by electrophoresis on ethidium bromide-stained, 1% agarose TBE gels and detected by trans-UV illumination using the Gel Doc XR+ (Bio-Rad) and Image Lab software (version 5.1). Detected bands were quantified using the Image Lab software (version 6.0.1). Results of the indicated band % per lane are shown as data points and mean.

#### Quantitative RT-PCR, Probe-based multiplex RT-PCR

Quantitative RT-PCR was performed with the GoTaq qPCR Master Mix (Promega) using 2% of cDNA in 10 μl reactions, 0.2 μM final concentration of sense and antisense primer (see Table S1 for sequences), and the CFX96 Touch Real-Time PCR Detection System (Bio-Rad) with Bio-Rad CFX Manager software (version 3.0). The reactions for each biological replicate were performed in triplicates and the Ct (threshold cycle) value was measured and average Ct values were calculated. For alternative splicing events, values for canonical isoforms were subtracted from values for NMD-sensitive isoforms to calculate the ΔCt. The mean log2 fold changes were calculated from three biologically independent experiments. Log2 fold change results are shown as data points and mean. Probe-based multiplex quantitative RT-PCRs were performed using the PrimeTime Gene Expression Master Mix (IDT) and the PrimeTime qPCR Assays containing primers and probes (IDT; ZFAS1 = Hs.PT.58.25163607, GAS5 = Hs.PT.58.24767969, B2M = Hs.PT.58v.18759587, TBP = Hs.PT.58v.39858774) following the manufacturer’s instructions. 2% of cDNA was used as a template in 10 μl reactions and samples were measured using CFX96 Touch Real-Time PCR Detection System (Bio Rad). The reactions for each biological replicate were performed in triplicates and the Ct (threshold cycle) value was measured and average Ct values were calculated. The Ct values of the housekeeping gene B2M or TBP (FAM-labelled) were subtracted from the target (ZFAS1, Cy5-labelled or GAS5, SUN-labelled) values to calculate the ΔCt. Three biologically independent experiments were used to calculate the mean log2 fold changes. The log2 fold changes are visualized as single data points and mean. All primers used in this study are listed in Table S1.

#### siRNA-mediated knock-downs

2.5-3.0 × 10^5^ cells were seeded in 6-well dish and reverse transfected using Lipofectamine RNAiMAX (Invitrogen) and 60 pmol siRNA following the manufacturer’s instructions. 48 after transfection cells were harvested in 1 ml in-house prepared TRI reagent ^60^ for RNA extraction or RIPA buffer (50 mM Tris/HCl pH 8.0, 0.1% SDS, 150 mM NaCl, 1% IGEPAL, 0.5% deoxycholate) for protein extraction. All siRNAs used in this study are listed in Table S1.

#### High-throughput-sequencing

The RNA was extracted and purified using the Directzol RNA MiniPrep kit including the recommended DNase I treatment (Zymo Research; Cat# R2052) according to manufacturer’s instructions. Libraries were prepared from 500 ng total RNA with the Illumina Stranded mRNA Preparation kit. ERCC RNA Spike-In Mix (Thermo Fisher) was added to the samples before library preparation. After poly-A selection (using Oligo(dT) magnetic beads), mRNA was purified, fragmented and reverse transcribed with random hexamer primers. Second strand synthesis with dUTPs was followed by A-tailing, adapter ligation and library amplification (12 cycles) to create the final cDNA libraries. After library validation and quantification (Agilent Tape Station), equimolar amounts of library were pooled. The pool was quantified by using the Peqlab KAPA Library Quantification Kit and the Applied Biosystems 7900HT Sequence Detection System. The pool was sequenced on an Illumina NovaSeq6000 sequencing instrument with a PE100 protocol aiming for 50 million clusters per sample. Following RNA-Seq datasets were obtained and analyzed: SMG5, SMG6, SMG7 knock-out/knock- down in HEK293 cells (BioStudies ^61,62^ accession E-MTAB-9330) ^7^, UPF1 degron (AID and dTAG/FKBP) in HEK293 or HCT116 cells (BioStudies accession E MTAB-13788, E-MTAB-13829) ^36^, UPF3A/B double knock-out in HEK293 cells (BioStudies accession E-MTAB-10716) ^40^.

#### Computational analyses of RNA-Seq data

For standard RNA-Seq analyses, reads were aligned against the human genome (GRCh38, GENCODE release 42 transcript annotations ^63^ supplemented with SIRVomeERCCome annotations from Lexogen; obtained from https://www.lexogen.com/sirvs/download/) using the STAR read aligner (version 2.7.10b) ^64^. Transcript abundance estimates were computed with Salmon (version 1.9.0) ^65^ in mapping-based mode using a decoy-aware transcriptome (GENCODE release 42) with the options –numGibbsSamples 30 –useVBOpt –gcBias –seqBias. After the import of transcript abundances in R (version 4.3.0) ^66^ using tximport (version 1.28.0) ^67^, differential gene expression (DGE) analysis was performed with the DESeq2 R package (version 1.40.1) ^68^. Genes with less than 10 counts in half the analyzed samples were pre-filtered and discarded. The DESeq2 log2FoldChange estimates were shrunk using the apeglm R package (version 1.22.1) ^69^. Differential transcript expression (DTE) analysis was performed using the Swish method from the fishpond R package (version 2.6.2) ^70^ based on 30 inferential replicate datasets drawn by Salmon using Gibbs sampling and imported via tximeta (version 1.18.1) ^71^. Transcripts were pre-filtered using 10 counts per transcript in at least one condition as cut-off. General significance cut-offs, as long as not indicated otherwise, were log2FoldChange > 1 & p.adjust < 0.0001 for DESeq2 DGE and log2FC > 1 & qvalue < 0.0001 for Swish DTE. Gene ontology functional enrichments analysis of gene lists, ordered by adjusted p-value, was performed using an ordered query by g:profiler via the R package gprofiler2 (version 0.2.2) ^72^, using gene ontology biological process (GO:BP) as data source, a list of all expressed/detected genes as custom background, domain scope set to ‘custom_annotated’ and with “fdr” multiple testing correction method applying significance threshold of 0.05. Most mRNA isoform properties were extracted from the GENCODE annotation and reference genome using R packages. Structure prediction was performed using RNAfold (version 2.6.4) ^73^.

#### Co-immunoprecipitation

Stable cell lines expressing FLAG-tagged SMG8 or endogenously tagged UPF1 were seeded in 10 cm dishes (2.5-3.0 x 10^6^ cells) and SMG8 expression was induced via Doxycycline. 2-3 days after seeding cells were harvested in 200 μl buffer E phos (20 mM HEPES-KOH (pH 7.9), 100 mM KCl, 10% glycerol, 1 mM DTT, Protease Inhibitor, 1x PhosSTOP (Roche), 1x Halt Protease and Phosphatase Inhibitor Cocktail (Thermo Scientific), 10 μg/μl RNase (Panreac AppliChem)). Cells were lysed using Bandelin Sonopuls mini20 with 15 pulses (2.5mm tip, 1 s pulse, 50% amplitude). Samples were adjusted to the same concentration and incubated for 2 h overhead shaking with Anti-FLAG M2 Magnetic Beads (Sigma-Aldrich). Beads were washed three times for5 min with mild wash buffer (20 mM HEPES-KOH (pH 7.9), 137 mM NaCl, 2 mM MgCl2, 0.2% Triton X-100, 0.1% NP-40). Co-immunoprecipitated proteins were eluted with SDS-sample buffer, separated by SDS-PAGE, and analyzed by immunoblotting.

#### Label-free quantitative mass spectrometry

Cells expressing endogenously FLAG-tagged UPF1 were seeded in 10 cm dishes (2.5-3.0 x 10^6^ cells). After 24 h, cells were treated with SMG1i ^39^ for 24 h and harvested in 200 μl buffer E phos (20 mM HEPES-KOH (pH 7.9), 100 mM KCl, 10% glycerol, 1 mM DTT, Protease Inhibitor, 1x PhosSTOP (Roche), 1x Halt Protease and Phosphatase Inhibitor Cocktail (Thermo Scientific), 10 μg/μl RNase (Panreac AppliChem)) and immunoprecipitation was performed as described above using mild wash buffer. Proteins were eluted using 44 μl FLAG-peptides (200 μg/ml; Merck/Sigma Aldrich) in 1x TBS. 44 μl of 10% SDS in 1x PBS was added and samples were incubated at 95 °C for 5 min. Samples were reduced with DTT at 55 °C for 30 min and alkylated with CAA at RT for 30 min (final concentrations 5 mM and 55 mM, respectively). Tryptic protein digestion was achieved by following a modified version of the single pot solid phase-enhanced sample preparation (SP3) ^74^. In brief, paramagnetic Sera-Mag speed beads (Thermo Fisher Scientific) were added to the reduced and alkylated protein samples and then mixed 1:1 with 100% acetonitrile (ACN). Protein-beads-complexes form during the 8-min incubation step, followed by capture using an in-house build magnetic rack. After two washing steps with 70% EtOH, the samples were washed once with 100% ACN. Then they were air-dried, resuspended in 5 μl 50 mM Triethylamonium bicarbonate supplemented with trypsin in an enzyme:substrate ratio of 1:50 and incubated for 16 h at 37°C. Afterwards, samples were acidified to 5% FA, and cleaned-up using SDB-RPS StageTips. Samples were loaded onto the tips, washed with 0.1% FA followed by washing with 80% AcN + 0.1% FA. Finally, samples were eluted with 40% NH_3_, dried down and resuspended in 4% AcN + 0.1% FA, ready for mass spectrometric analysis.

#### Data Acquisition

Samples were analyzed by the CECAD Proteomics Facility on an Orbitrap Exploris 480 (Thermo Scientific, granted by the German Research Foundation under INST 216/1163-1 FUGG) mass spectrometer equipped with a FAIMSpro differential ion mobility device that was coupled to an Vanquish neo in trap-and-elute setup (Thermo Scientific). Samples were loaded onto a precolumn (Acclaim 5μm PepMap 300 μ Cartridge) with a flow of 60 μl/min before reverse-flushed onto an in-house packed analytical column (30 cm length, 75 μm inner diameter, filled with 2.7 μm Poroshell EC120 C18, Agilent). Peptides were chromatographically separated with an initial flow rate of 400 nl/min and the following gradient: initial 2% B (0.1% formic acid in 80% acetonitrile), up to 6% in 3 min. Then, flow was reduced to 300 nl/min and B increased to 20% B in 26 min, up to 35% B within 15 min and up to 98% solvent B within 1.0 min while again increasing the flow to 400 nl/min, followed by column wash with 95% solvent B and reequilibration to initial condition. The FAIMS pro was operated at -40 V compensation voltage and electrode temperatures of 99.5 °C for the inner and 85 °C for the outer electrode. The mass spectrometer was operated in data-dependent acquisition top 24 mode with MS1 scans acquired from 350 m/z to 1400 m/z at 60k resolution and an AGC target of 300%. MS2 scans were acquired at 15k resolution with a maximum injection time of 22 ms and an AGC target of 300% in a 1.4 Th window and a fixed first mass of 110 m/z. All MS1 scans were stored as profile, all MS2 scans as centroid.

#### Sample Processing in MaxQuant

All mass spectrometric raw data were processed with MaxQuant (version 2.4) ^75^ using default parameters against the Uniprot Human canonical reference proteome database (UP5640) with the match-between-runs option enabled between replicates. Label-free quantification was performed separately for replicate group to better cope with strong differences in protein abundances in IP situations. Follow-up analysis was done in Perseus 1.6.15^76^. Protein groups were filtered for potential contaminants and insecure identifications. Remaining IDs were filtered for data completeness in at least one group and missing values imputed by sigma downshift (0.3 σ width, 1.8 σ downshift). Afterwards, FDR-controlled two-sided t-tests were performed. Finally, majority protein IDs were used for protein annotations and further analysis.

#### Data presentation

Schematic representations and figures were prepared/assembled using CorelDraw 2017. Quantifications and calculations for other experiments were performed - if not indicated otherwise - with Microsoft Excel (version 1808 or 2311) or R (version 4.3.0) and all plots were generated using IGV (version 2.14.1) ^77^, GraphPad Prism 5, ggplot2 (version 3.4.2) ^78^, ggsashimi (version 1.1.5) ^79^, nVennR (version 0.2.3) ^80^ or ComplexHeatmap (version 2.18.0) ^81^. If not indicated otherwise, the box of boxplots extends to the 25th and 75th percentile with the median in bold line, outliers are not shown.

#### Quantification and statistical analysis

Most performed statistical tests are already implemented in the used bioinformatic tools. For differential gene expression (DGE) analysis, p-values were calculated by DESeq2 using a two-sided Wald test and corrected for multiple testing using the Benjamini-Hochberg method. For differential transcript expression (DTE) analysis, p-values were calculated by Swish using a Mann–Whitney Wilcoxon test on inferential replicate count matrices and corrected for multiple testing using q-value approaches. For ECDF plots, two-sided Kolmogorov-Smirnov tests were performed using the stats R package. Linear regression of scatter plots was performed using the stat_poly_eq function of the ggpmisc R package, displaying the adjusted coefficient of determination.

## Supplemental Information

**Figure S1.**
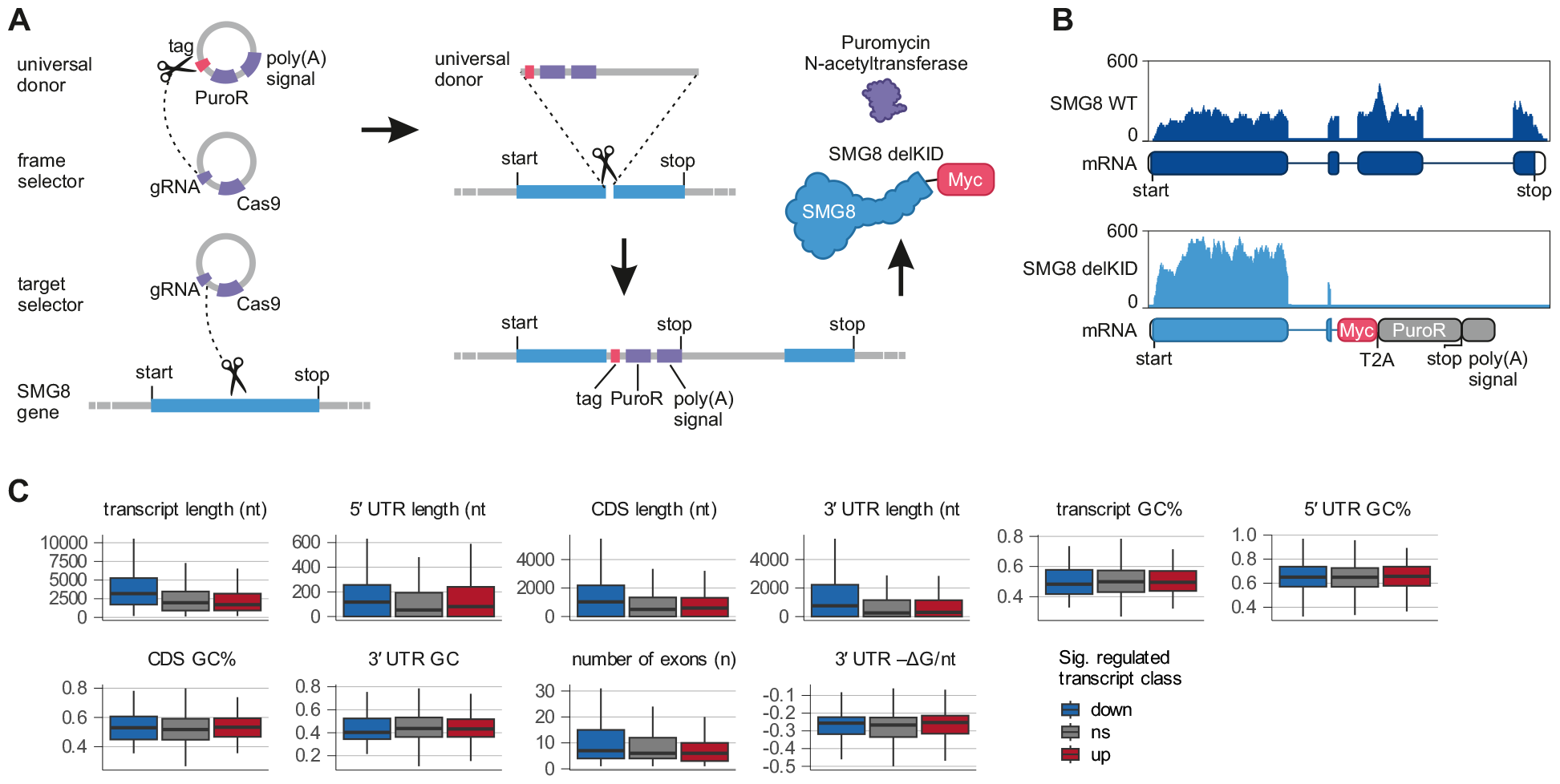
Overview of the CRISPaint system and further analyses of SMG8 delKID RNA-Seq data. (**A**) Schematic overview of the generation of SMG8 delKID clone via the CRISPaint system ^32^. The universal donor encodes for a Myc tag, the Puromycin resistance gene (PuroR) and contains a poly(A) signal. The frame selector encodes for Cas9 and a gRNA targeting the universal donor. The target selector encodes for Cas9 and a gRNA targeting the SMG8 gene. The linearized universal donor is inserted into the SMG8 gene resulting in the expression of truncated SMG8 and Puromycin N-acetyltransferase. (**B**) Read coverage of SMG8 gene from WT and SMG8 delKID RNA-Seq data is shown as Integrative Genomics Viewer (IGV) snapshots. Y-axis (reads) was scaled equally for both conditions and SMG8 WT transcript and the modified SMG8 delKID transcript is shown below. (**C**) Boxplot of annotation-derived transcript properties or 3’ UTR length-normalized minimum thermodynamic free energy (–ΔG/nt) of significantly regulated transcripts (down: blue, up: red) or not significantly changed transcripts (ns) in SMG8 delKID cells.

**Figure S2.**
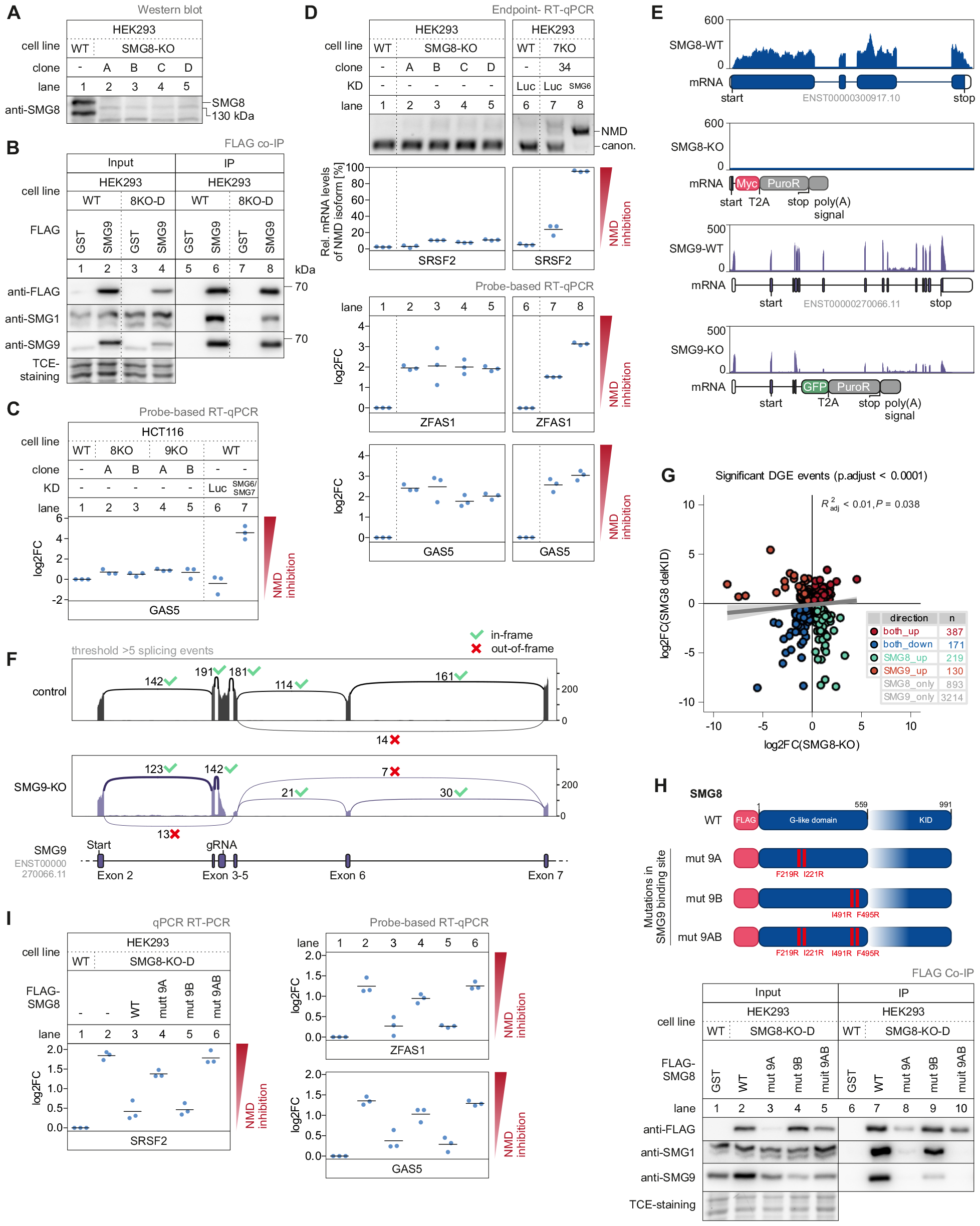
Characterization of SMG8- and SMG9-KO cells. (**A**) Western blot analysis of different HEK293 SMG8-KO cell lines (clone A, B, C, D) using anti-SMG8 antibody (AK-159; n=1 biologically independent sample; see Table S1 for antibody details). (**B**) Western blot after FLAG co-immunoprecipitation (IP) of FLAG-tagged GST (control) or SMG9 constructs in WT or SMG8-KO (clone D) cells. Anti-FLAG (AK-115), anti-SMG1 (AK-088) and anti-SMG9 (AK-170) antibodies were used and TCE-staining serves as a control (n=3 biologically independent samples). (**C**) Quantitative probe-based RT-PCR of GAS5 in WT, SMG8-KO or SMG9-KO cells with or without indicated knock- down (KD). The ratio of GAS5 to the TBP reference was calculated; data points and means from the qPCRs are plotted as log2 fold change (log2FC) (n=3 biologically independent samples). (**D**) End-point RT-PCR detection of SRSF2 transcript isoforms (top) and quantitative probe-based RT-PCR (bottom) of ZFAS1 and GAS5 in WT or SMG8-KO cells with or without indicated KD. The detected SRSF2 isoforms are indicated on the right (NMD = NMD-inducing isoform; canon. = canonical isoform). Relative mRNA levels of SRSF2 isoforms were quantified from bands of agarose gels (n=3 biologically independent samples). The ratio of ZFAS1 or GAS5 to the TBP reference was calculated; data points and means from the qPCRs are plotted as log2 fold change (log2FC) (n=3 biologically independent samples). (**E**) Read coverage of SMG8/SMG9 gene from SMG8/SMG9 WT and SMG8/SMG9-KO RNA-Seq data is shown as Integrative Genomics Viewer (IGV) snapshots. Y-axis (reads) was scaled equally for both conditions and SMG8/SMG9 WT transcript and the modified SMG8/SMG9-KO transcript is shown below. (**F**) Sashimi plot of SMG9 read coverage and junction-spanning read counts from SMG9 WT and SMG9-KO RNA-Seq data. Y-axis (reads) was scaled equally for both conditions and SMG9 transcript (exon 2-7) is depicted below. Cutoff for junction reads was set to >5 and splicing events leading to in-frame (green tick) or out-of-frame (red cross) open reading frames are indicated. (**G**) Scatter plot of differentially regulated genes in SMG8 delKID against SMG8-KO cells. Changes in gene expression are shown as log2FC (p.adjust < 0.0001). Linear regression with p-value (P) and adjusted coefficient of determination is shown. (**H**) Schematic representation of the SMG8 domain structure. The mutated constructs with impaired SMG9 interaction are shown (mut 9A, 9B, 9AB). Western blot after FLAG co-immunoprecipitation (IP) of FLAG-tagged GST (control) or SMG8 constructs in WT or SMG8-KO cells is shown. Anti-FLAG (AK-115), anti-SMG1 (AK-088) and anti-SMG9 (AK-170) antibodies were used. TCE-staining serves as a control (n=3 biologically independent samples). (**I**) End-point RT-PCR detection of SRSF2 transcript isoforms (top) and quantitative probe-based RT-PCR (bottom) of ZFAS1 and GAS5 in WT or SMG8-KO cells upon expression of the indicated FLAG-tagged rescue constructs. The detected SRSF2 isoforms are indicated on the right (NMD = NMD-inducing isoform; canon. = canonical isoform). Relative mRNA levels of SRSF2 isoforms were quantified from bands of agarose gels (n=3 biologically independent samples). The ratio of ZFAS1 or GAS5 to the B2M reference was calculated; data points and means from the qPCRs are plotted as log2 fold change (log2FC) (n=3 biologically independent samples).

**Figure S3.**
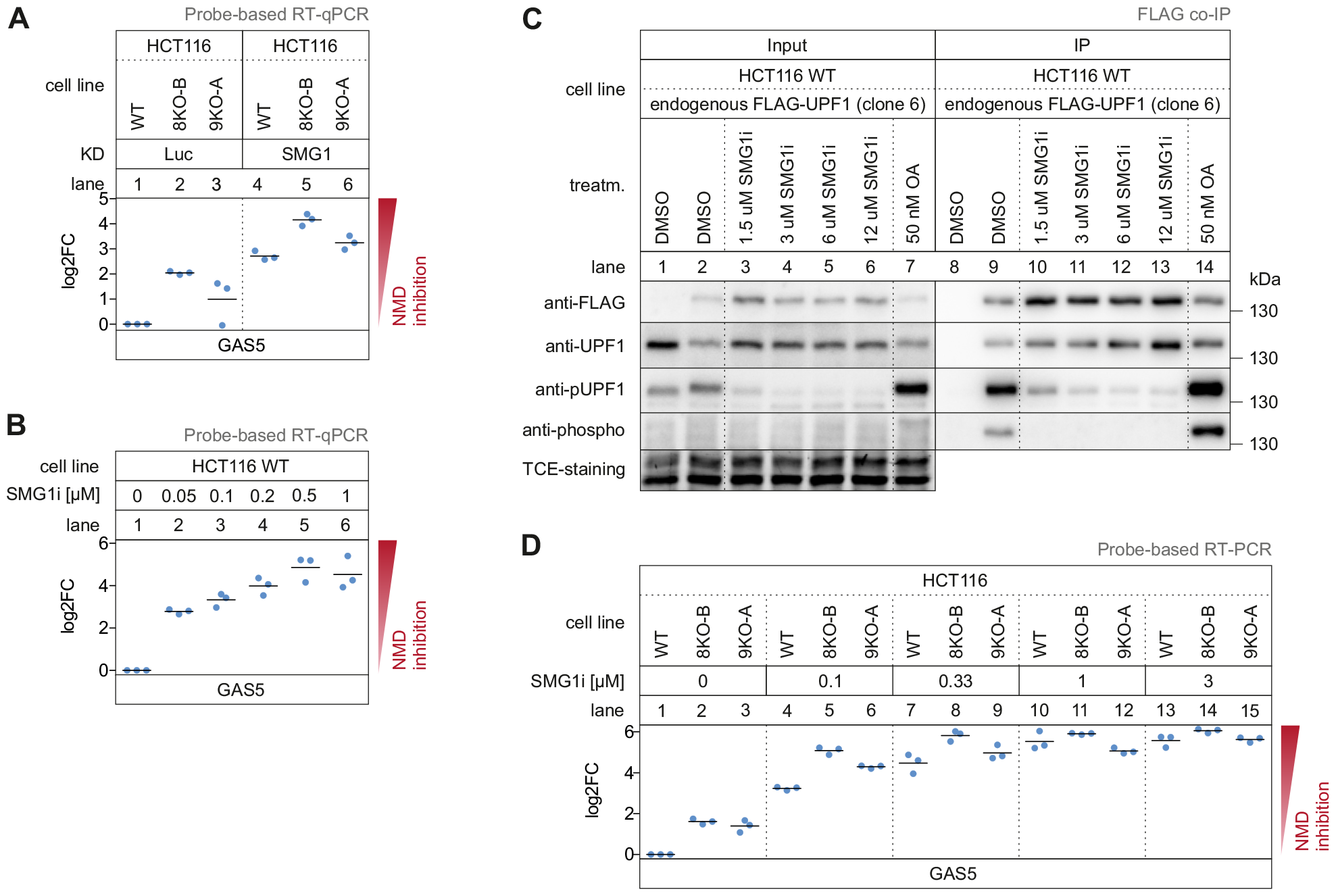
SMG1i treatment of WT, SMG8-KO and SMG9-KO cells. (**A**) Quantitative probe-based RT-PCR of GAS5 in WT, SMG8-KO or SMG9-KO cells with Luc or SMG1 knock-down (KD). The ratio of GAS5 to the B2M reference was calculated; data points and means from the qPCRs are plotted as log2 fold change (log2FC) (n=3 biologically independent samples). (**B, D**) Quantitative probe-based RT-PCR of GAS5 in WT cells (B), SMG8-KO and SMG9-KO cells (D) with treatment of different SMG1i concentrations for 24 h. The ratio of GAS5 to the B2M reference was calculated; data points and means from the qPCRs are plotted as log2 fold change (log2FC) (n=3 biologically independent samples). (**C**) Western blot after FLAG co-immunoprecipitation (IP) of untagged (control) or endogenously FLAG-tagged UPF1 in WT cells treated with indicated concentrations of SMG1i for 24 h or Okadaic acid for 2 h. Anti-FLAG (AK-115), anti-UPF1 (AK-156), anti-pUPF1 (serine 1116; AK-146) and anti-phospho (binds phosphorylated serine/threonine; AK-126) antibodies were used. TCE-staining serves as a control (n=2 biologically independent samples; see Table S1 for antibody details).

**Figure S4.**
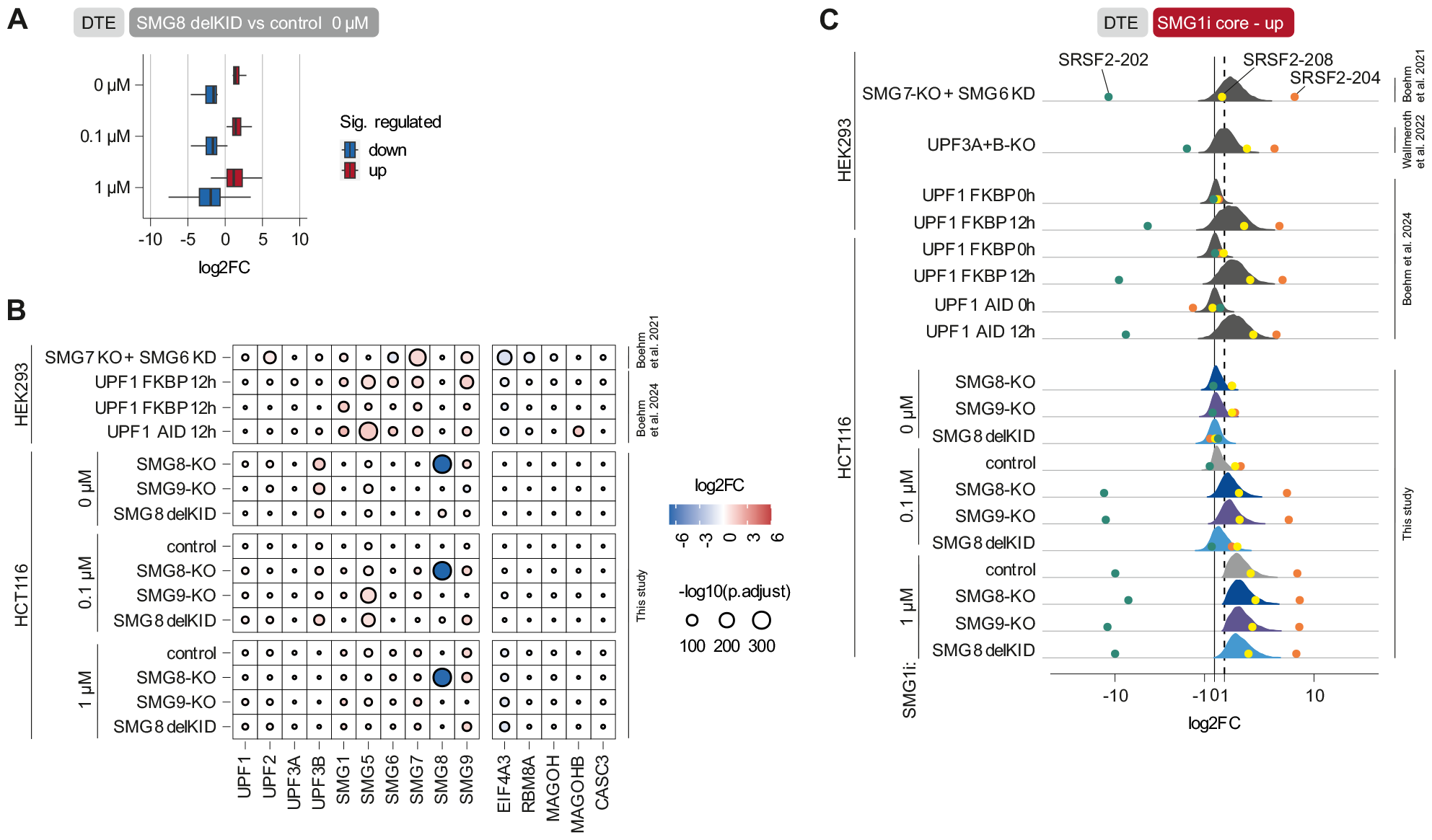
RNA-Seq analysis of SMG1i-treated WT, SMG8-KO and SMG9-KO cells. (**A**) Differentially up- or downregulated genes (p.adjust < 0.0001 & |log2FC| > 1) in SMG8 delKID cells with 0 μM SMG1i are compared to SMG8 delKID cells with different concentrations of SMG1i. (**B**) RNA-Seq data of WT, SMG8-KO, SMG9-KO and SMG8 delKID cells treated with different concentrations of SMG1i for 24 h were compared with SMG7-KO + SMG6 knock-down (KD; clone 34) ^7^ or three UPF1 degron conditions ^36^ regarding the expression changes of NMD and EJC factors. (**C**) Distribution of expression changes of the upregulated SMG1i core transcripts in RNA-Seq data obtained in this study and compared to SMG7- KO + SMG6-KD (clone 34) ^7^, UPF3A/B-KO ^40^ and three UPF1 degron conditions ^36^. Canonical (green) and NMD-annotated isoforms (yellow, orange) of SRSF2 are indicated as circles.

**Figure S5.**
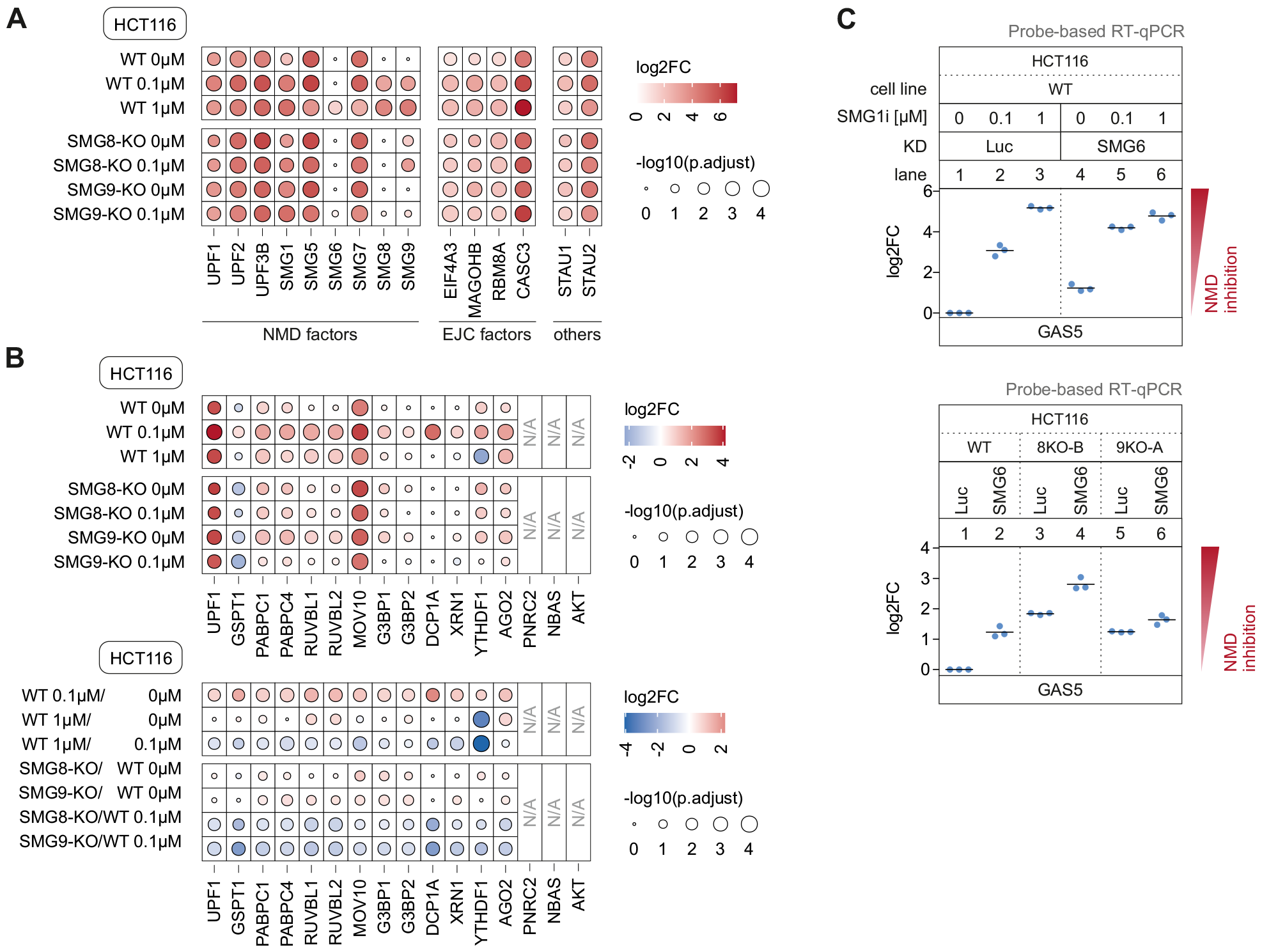
Proteomic analysis of UPF1 interactome in WT, SMG8-KO and SMG9-KO cells upon SMG1i treatment. (**A, B**) Heatmap of mass spectrometry-based analysis of FLAG co-immunoprecipitated (IP), untagged (control) or endogenously FLAG-tagged UPF1 in WT, SMG8-KO and SMG9-KO cells. Cells were treated with indicated concentrations of SMG1i for 24 h. Colored points indicate the log2 fold change (log2FC) and point size corresponds to the adjusted p-value (adj. p-value; from Students t test; n = 4 biologically independent samples). (**C**) Quantitative probe-based RT-PCR of GAS5 in WT, SMG8-KO and SMG9-KO cells with Luc knock-down (KD; control) or SMG6-KD. WT cells were treated in addition with SMG1i for 24 h. The ratio of GAS5 to the B2M reference was calculated; data points and means from the qPCRs are plotted as log2 fold change (log2FC) (n=3 biologically independent samples).

## Supplemental Tables

Table S1. Resource and material lists

Table S2. Differential gene and transcript expression analysis

Table S3. Core target definitions

Table S4. Proteomic analyses of UPF1 interactome

